# Single-stranded telomeric repeats segregate into spatial compartments within ALT-associated PML bodies

**DOI:** 10.64898/2026.05.10.724127

**Authors:** Emma S. Koeleman, Charlotte Kaplan, Maria Augusta R. B. F. Lima, Delia M. Braun, Caroline Knotz, Karsten Rippe

## Abstract

Alternative lengthening of telomeres (ALT)-associated PML bodies (APBs) concentrate single-stranded (ss) telomeric DNA and RNA species critical for recombination-based telomere maintenance. However, how these species are organized inside APBs has remained invisible at microscopic resolution. Here, we map the nanoscale topology of APB components using 3D MINFLUX super-resolution microscopy combined with multiplexed exchange DNA-PAINT labeling at ∼3 nm localization precision. We discover that ssC-rich and ssG-rich telomeric repeats occupy distinct spatial compartments within a partially open, ∼70 nm-thick PML protein shell: ssC-rich repeats concentrate at the inner shell surface, while ssG-rich repeats distribute broadly throughout the interior alongside TRF1-marked double-stranded telomeric chromatin. The ssG-rich signal is predominantly DNA and frequently colocalizes with POT1 assemblies. Additional TRF1 clusters outside the shell indicate multi-telomere association. Together, the spatial segregation of single-stranded telomeric species within APBs imposes structural constraints on different mechanisms of ALT-mediated telomere elongation and defines features that distinguish them.

**Graphical abstract:** 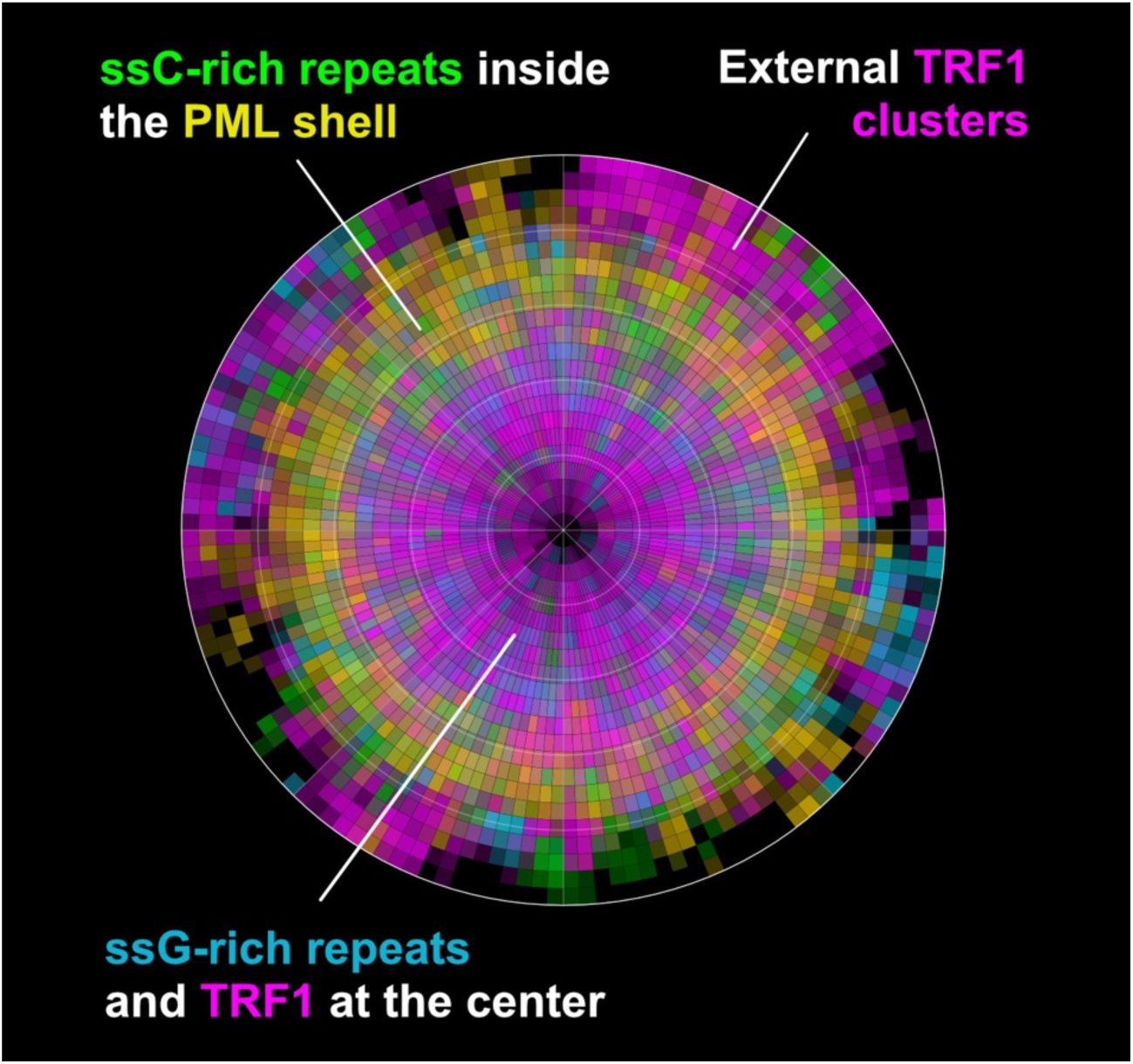

**Highlights:**

- 3D MINFLUX combined with exchange DNA-PAINT resolves APB components at ∼3 nm isotropic precision, enabling direct visualization of molecular organization beyond the reach of conventional super-resolution microscopy.
- PML forms a partially open, ∼70 nm thick spherical shell in APBs with reduced molecular density compared to canonical PML nuclear bodies, providing the structural scaffold and coordinate system for mapping internal APB organizationa.
- Double-stranded telomeric chromatin (TRF1-marked) fills the APB interior, with additional TRF1 domains outside the shell indicative of multi-telomere clustering at individual APBs.
- ssC-rich repeats concentrate in the inner part of the PML shell while ssG-rich DNA repeats distribute broadly through the interior, revealing strand-specific spatial segregation.
- The ssG-rich single-stranded repeats in APBs predominantly comprise DNA and partially colocalize with POT1, connecting their spatial distribution to single-stranded telomeric DNA substrates.
- The spatial organization of APB components motivates a mechanistic model in which t-loop processing generates C-circle templates for rolling-circle amplification, while multi-telomere clustering facilitates inter-telomeric recombination.

## Introduction

Unlimited proliferation of cancer cells requires activation of a telomere maintenance mechanism, either via telomerase or the alternative lengthening of telomeres (ALT) pathway ^1,2^. ALT is used by approximately 15% of cancer entities ^3^ and exploits aberrant DNA repair and recombination pathways to elongate telomeric repeats ^4,5^. Its hallmark is the formation of promyelocytic leukemia nuclear bodies (PML-NBs; 0.2-1µm in diameter) at certain telomeres, known as ALT-associated PML nuclear bodies (APBs) ^6^. In APBs, PML proteins assemble around a core of shelterin proteins and telomeric repeats ^7^, forming a structure enriched for single-stranded (ss) C- and G-rich telomeric repeats, detectable by native FISH ^8,9^. ALT activity is reflected by the production of C-circles, which are predominantly single-stranded telomeric DNA circles composed of C-strand repeats (CCCTAA)_n_ generated during telomere processing ^10,11^. In addition, C- and G-rich single-stranded telomeric DNA overhangs and telomere-internal regions are present ^12^. The telomere repeat overhangs can invade the duplex telomeric repeat tract to form a telomere loop (t-loop), a structure visualized in vitro by electron microscopy and direct stochastic optical reconstruction microscopy (dSTORM) ^13,14^. Finally, telomeric repeat-containing RNA (TERRA), a long noncoding RNA transcribed from subtelomeric promoters into UUAGGG repeats, associates with telomeres via RNA:DNA hybrid (R-loop) formation, further expanding the repertoire of single-stranded telomeric substrates present at APBs ^15–17^. Together, these single-stranded telomeric repeat sequences accumulate within APBs and are thought to drive recombination-based telomere elongation in ALT.

The diverse DNA and RNA repeat structures underscore the molecular complexity of APBs and imply a high degree of spatial and functional organization crucial to the ALT mechanism. However, the relative nanoscale arrangement of PML, double-stranded telomeric chromatin, and strand-specific single-stranded telomeric repeats within APBs has remained inaccessible because of microscopy resolution limits. As a consequence, key mechanistic questions remain unresolved, including whether distinct telomeric substrates (dsDNA, ssG-rich DNA/RNA, ssC-rich DNA) are spatially segregated within APBs, how many telomeres are engaged per APB, and how APB architecture facilitates strand invasion, recombination, or rolling-circle–based telomere synthesis ^11,18,19^. Bridging this spatial knowledge gap requires direct visualization of APB nanoarchitecture at molecular length scales, which has so far remained out of reach.

A new venue for acquiring this information is MINFLUX (minimal photon fluxes) super-resolution fluorescence microscopy. It employs a doughnut-shaped excitation beam repositioned around a fluorophore at targeted coordinates, and the fluorophore’s position is estimated from the relative photon counts recorded at each beam position. Iteratively reducing the scan diameter around the targeted coordinates during acquisition progressively refines this estimate, achieving isotropic localization precision down to 2 nm ^20^. This nanometer-scale localization precision has enabled the characterization of diverse cellular structures, including mitochondrial protein complexes, membrane protein conformational states, and nucleosomes^21–23^.

However, MINFLUX has not yet been used to resolve the molecular topology of protein-nucleic acid assemblies within chromatin compartments and condensates at the 0.1-1 µm mesoscale, which play an important role in enriching and targeting genome-associated activities ^24–26^. Here, we apply it to resolve the nanoscale three-dimensional (3D) topology of telomeric repeats and the shelterin components TRF1 and POT1 within APBs. We used TRF1 as a marker for telomeric dsDNA and POT1 to identify ssG-rich DNA ^27,28^. POT1 forms a heterodimer with TPP1 and binds single-stranded G-rich telomeric DNA overhangs, with TIN2, another shelterin protein, linking TRF1 and TRF2 dimers to POT1-TPP1 ^29^. With this approach, we discover that ssC-rich and ssG-rich telomeric repeats are not randomly intermingled but occupy distinct spatial compartments within APBs. This strand-specific spatial segregation, together with the architecture of the PML shell and multi-telomere clustering, defines structural constraints that any mechanistic model of ALT must accommodate.

## Results

### 3D MINFLUX microscopy with exchange DNA-PAINT enables nanoscale mapping of APB architecture

We combined 3D MINFLUX with exchange DNA-PAINT ^30,31^ and automated fluidics to image three targets within individual APBs: PML, TRF1 and single-stranded telomeric repeats (**Fig. 1A**). Sequential exchange of Cy3B-conjugated imager strands during continuous MINFLUX measurements enabled multiplexed labeling in a single experiment (see Methods). During imaging, MINFLUX records the three-dimensional position of individual fluorophores as x, y, z coordinates, referred to as localizations. Multiple localizations can originate from the same fluorophore, linked by a shared identifier, and each carries a quality parameter that enables filtering and calculation of localization precision (see Methods). For each target, we achieved a median lateral localization precision of σ_x,y_ ∼ 3.7-4.4 nm and σ_z_ ∼ 2.5 nm (**Fig. 1B, C**). APBs were identified in U2OS cells stably expressing endogenous HaloTag-PML and inducible TRF1-GFP (**Supplementary Fig. 1**), enabling identification of co-localization prior to DNA-PAINT imaging (**Fig. 1D**). Representative 3D MINFLUX datasets are shown for each target (**Fig. 1E, Supplementary Video 1-4**). This combination of isotropic nanometer-scale precision and multiplexed target detection enabled us to resolve the spatial relationships between PML, TRF1 and single-stranded telomeric repeats within individual APBs.

**Fig. 1.**
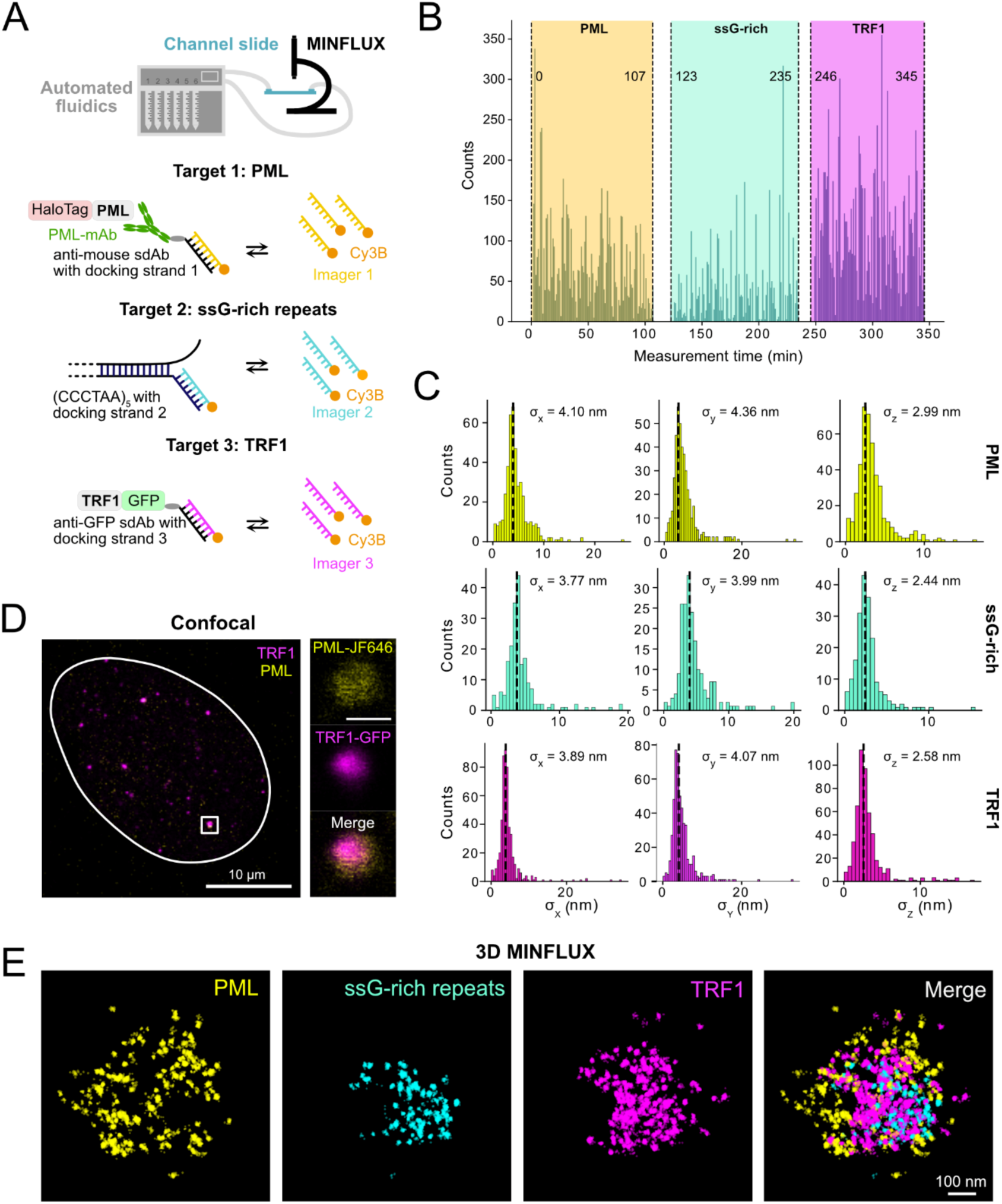
3D MINFLUX microscopy combined with exchange DNA-PAINT enables nanometer-scale localization of APB components. (**A**) Scheme of the multi-target exchange DNA-PAINT approach for MINFLUX imaging. Three targets were visualized: PML (detected via anti-mouse antibody linked to single-domain antibody with docking strand 1), ssG-rich telomeric repeats (detected via (CCCTAA)₅ probe with docking strand 2) and TRF1-GFP (detected via anti-GFP single-domain antibody with docking strand 3). Sequential automated fluidics exchange between imager strands 1, 2 and 3 conjugated to a Cy3B fluorophore enabled multiplex detection of the three targets. (**B**) Representative time-resolved visualization of a 3-target imaging acquisition, showing sequential detection of individual targets and temporal transitions between imaging rounds. (**C**) Localization precision of all three targets in x, y, and z dimensions. Histograms show the distribution of localization precision values, with median per-trace values of 3.7 to 4.4 nm in x and y and 2.5 nm in z. The vertical dashed lines indicate the median per-trace localization precision. (**D**) Confocal imaging showing colocalization of HaloTag-PML (detected with Janelia Fluor 646 ligand) and TRF1-GFP in U2OS ALT⁺ cells to identify APB structures. Scale bar, 10 μm (main) and 500 nm (inset). (**E**) 3D MINFLUX visualization of an APB showing the spatial organization of PML (yellow), TRF1 (magenta), and ssG-rich repeats (cyan). See also Supplementary Video 1-4. Scale bar, 100 nm.

### PML assembles into a partially open spherical shell with reduced density in APBs compared to PML-NBs

To characterize the nanoscale architecture of the PML shell, we applied density-based spatial clustering of applications with noise (DBSCAN) to identify individual PML structures (see Methods) (**Fig. 2A**). To assess whether APBs differ structurally from canonical PML-NBs lacking telomeres, we fitted a sphere to each PML cluster and found that both types assemble into a spherical shell with an inhomogeneous molecular distribution (**Fig. 2B**). To quantify potential differences, we compared shell radius, thickness, molecule number, and density across all structures (**Fig. 2C**), detecting between 50 and 700 PML localizations per structure. Although mean shell thickness was identical at 68 nm, APBs had a larger mean radius (192 nm versus 130 nm in PML-NBs) and a lower volume-normalized PML density, indicating a less compact nanoscale organization (**Fig. 2C**).

**Fig. 2.**
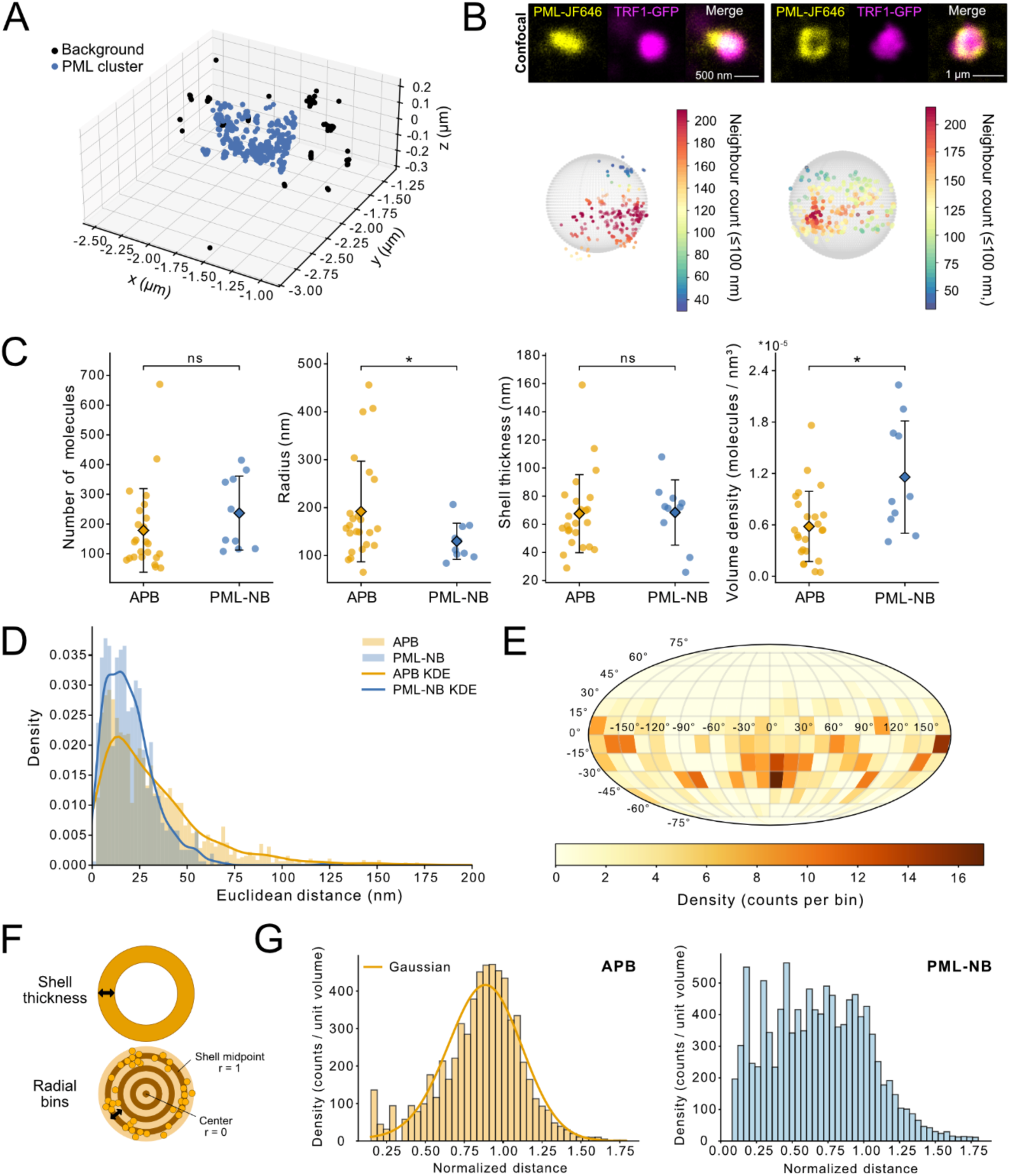
Nanoscale topology of PML in APBs compared to PML-NBs. (**A**) DBSCAN clustering of one APB acquisition. All localizations outside the APB cluster were excluded from further analysis. (**B**) Representative confocal images of two APBs showing colocalization of HaloTag-PML detected with Janelia Fluor 646 (yellow) and TRF1-GFP (magenta). Scale bars, 500 nm (left) and 1 μm (right). Corresponding 3D MINFLUX reconstruction of PML shell architecture after DBSCAN clustering (bottom panel) with fitted sphere overlay (gray wireframe). The color scale indicates the number of neighboring molecules within 100 nm along the spherical surface. (**C**) Point plots of quantitative parameters for APBs (n=24) compared to PML-NBs (n=10): number of molecules, shell thickness, radius and volume density (mean ± SD and mean values). Points represent individual structures; diamonds show the mean. Statistical significance between groups was assessed using a two-sided Welch’s t-test. (**D**) First nearest-neighbor (NN) distance distributions for APBs (orange, n=24) and PML-NBs (blue, n=10), shown as histograms with kernel density estimates (KDE) overlaid. APB and PML-NB distributions differ significantly (Kolmogorov–Smirnov test: D = 0.240, p = 6.6·10^−30^). (**E**) Heatmap of PML molecule density across the shell surface of a representative APB, with localizations binned into 24 horizontal and 12 vertical bins. Color scale indicates density in counts per bin. (**F**) Schematic illustration of the radial binning approach used to quantify PML molecule density across the shell depth. Molecules are counted within concentric rings of fixed width and variable radius, and counts are normalized to the volume of each bin to obtain a volume-corrected density profile. (**G**) Volume-normalized PML molecule density distribution across all APBs. Normalized distance of each PML molecule to the center was quantified for radial bins, which were normalized to shell volume per bin. A Gaussian fit was applied to the data (μ = 0.997 and σ = 0.222) (n=24). (**H**) Volume-normalized PML molecule density distribution across all PML-NBs. Normalized distance of each PML molecule to the center was quantified for radial bins which were normalized to shell volume per bin (n=10). The NB distribution could not be well described by a Gaussian model.

To probe molecular spacing directly, we compared nearest-neighbor (NN) Euclidean distances, which were larger in APBs (Kolmogorov–Smirnov test, D = 0.24, p = 6.6·10^−30^) (**Fig. 2D**, **Supplementary Fig. 2**), consistent with the high variability in local PML density across the shell illustrated for a representative structure (**Fig. 2E**). To determine whether APBs and PML-NBs differ in how PML molecules are distributed across the shell depth, we analyzed volume-corrected radial localization frequencies, with positions normalized from the structure center (r = 0) to the PML shell (r = 1). APBs exhibited a Gaussian distribution centered on the shell midpoint, whereas PML-NBs showed a more uniform distribution across the shell (**Fig. 2F, G**). Together, these results show that both PML-NBs and APBs form partially open spherical shells. APBs are less densely packed and display a Gaussian distribution of molecules across the shell depth.

### TRF1 clusters inside APBs and forms discrete external domains consistent with multi-telomere association

Next, we evaluated the spatial organization of TRF1 as a marker for double-stranded telomeric repeats in APBs. TRF1 molecules were predominantly located inside the APB, with an average of 812 molecules, and a smaller subset with an average of 107 molecules outside (**Fig. 3A, B**). TRF1 structure size was positively correlated with the radius of the PML shell (ρ = 0.58; **Supplementary Fig. 4C**), consistent with coordinated scaling of TRF1 assemblies and PML structures, in agreement with previous reports ^32^. Since TRF1 scaling with PML radius suggested structural integration of double-stranded telomeric DNA within the APB, we next asked whether single-stranded telomeric repeats showed a similar relationship. Interestingly, ssG-rich repeats showed a similar correlation (ρ = 0.70), whereas ssC-rich repeats were less strongly associated (ρ = 0.32) (**Supplementary Fig. 4C**). We next probed the nanoscale organization of TRF1 molecules within APBs using NN analysis. TRF1-TRF1 distances showed a kernel density estimation (KDE) peak at 12.7 nm, whereas the first NN distance to PML peaked at 40.8 nm (**Fig. 3C**). The short TRF1–TRF1 distances are consistent with the predicted spacing of TRF1-GFP dimers (**Supplementary Fig. 4B**). Improving the mean localization precision through photon aggregation did not alter the observed NN distances (**Supplementary Fig. 3, 4A**).

**Fig. 3.**
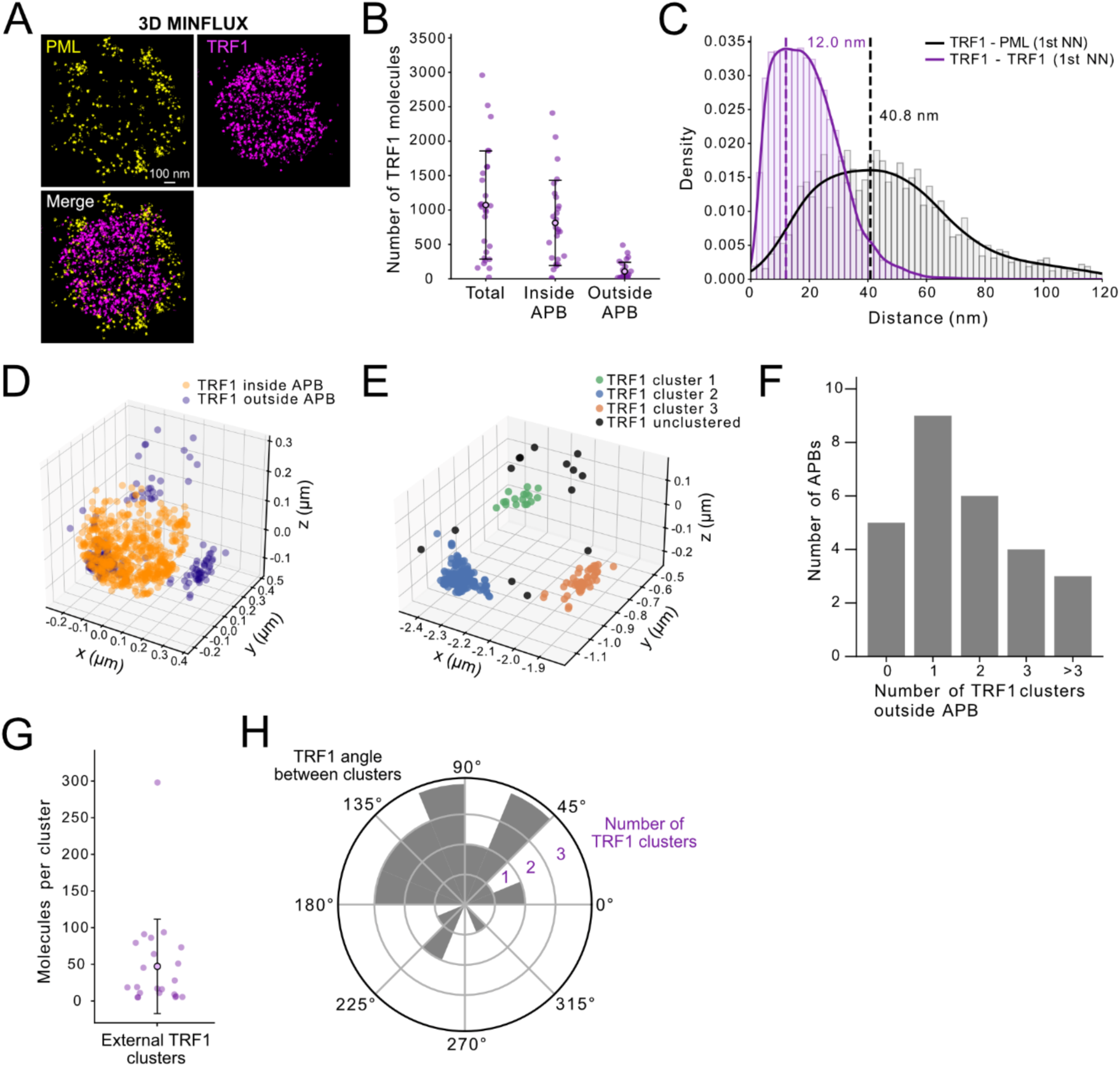
Nanoscale organization of TRF1 reveals multi-telomere association within APBs. (**A**) Representative 3D MINFLUX visualization of the TRF1 distribution (magenta) relative to the PML shell (yellow). Scale bar, 100 nm. (**B**) Quantification of TRF1 molecule count per APB structure, showing total number of TRF1 molecules, molecules inside APB, and molecules outside APB (n=27). Point plots display mean ± SD with individual data points. (**C**) First nearest-neighbor (NN) distance distributions for TRF1–TRF1 (purple) and TRF1–PML (black) pairs, shown as histograms with KDE curves. Dashed lines indicate KDE peaks at 12.7 nm and 40.8 nm, respectively. (**D**) Visualization of TRF1 molecules localizing inside (orange) or outside (purple) the APB structure. (**E**) DBSCAN clustering of external TRF1 molecules. TRF1 molecules are assigned to discrete clusters (green, orange and blue) or assigned as unclustered (black). (**F**) DBSCAN clustering analysis of TRF1 molecules outside APBs, revealing multiple distinct clusters. Distribution shows number of structures with 1, 2, 3, or >3 external TRF1 clusters (n=27). (**G**) Quantification of the size of external TRF1 clusters. Point plots display mean ± SD with individual data points. (**H**) Angular distribution analysis of external TRF1 clusters. Only APBs with more than 1 external cluster were included. For each structure, the TRF1 cluster with the highest molecule count was set at 0° and the angle to each next cluster was computed (n=27). Angles were assigned to 22.5° bins.

To assess the spatial distribution of TRF1 relative to the PML scaffold, we mapped TRF1 onto the normalized radius of the PML shell, classifying TRF1 molecules as inside (r ≤ 1) or outside (r > 1) the APB (**Fig. 3D, Supplementary Fig. 4D**). In 19% of APBs, all TRF1 clusters were confined within the PML shell, whereas in the remaining 81%, we detected TRF1 clusters outside the fitted shell boundary. Clustering analysis of external TRF1 localizations identified spatially discrete clusters with an average size of 47 molecules (**Fig. 3E, G**).

APBs contained on average 1.7 external clusters with the majority displaying one or two external clusters (**Fig. 3F**). The presence of multiple spatially separated TRF1 clusters outside the PML shell is consistent with the association of multiple telomeres within individual APBs^33^. Angular analysis of external TRF1 clusters revealed no preferred orientation, with a mean angular spacing of 40 degrees, indicating non-preferential rather than ordered clustering geometry (**Fig. 3H**). Together, these observations indicate that individual APBs are associated with multiple telomeres without a fixed geometric arrangement.

### ssC-rich repeats enrich at the PML shell while ssG-rich repeats distribute across the APB interior

Having established the PML shell as the structural scaffold and TRF1-marked chromatin as the core component, we next examined how the different single-stranded telomeric repeat species are organized within this framework. To quantify spatial organization, we analyzed the volume-corrected radial distribution of localizations by normalizing distances from the APB center (r = 0) to the shell boundary (r = 1). For each component, a center of mass (CoM) was calculated based on the radial distribution of localizations. TRF1 localized predominantly toward the middle of the APB (CoM = 0.49), whereas ssC-rich repeats localized toward the shell (CoM = 0.72) (**Fig. 4A, B, Supplementary Video 5-8**). Despite this peripheral enrichment, the majority of ssC-rich molecules were located within the APB, suggesting that the shell provides some barrier for single-stranded repeats (**Fig. 4F**). In contrast, ssG-rich repeats were more broadly distributed across the APB (CoM = 0.57; **Fig. 4C, D**). Permutation testing confirmed that the radial distributions of ssG-rich and ssC-rich repeats differ strongly (p = 0.0056) (**Fig. 4E**). Together, these results demonstrate that ssC-rich repeats accumulate in the inner part of the PML shell while ssG-rich repeats are broadly distributed across the APB interior. This broader distribution may reflect the detection of both DNA and RNA by the ssG-rich probe, suggesting a potential contribution of RNA to the observed signal.

**Fig. 4.**
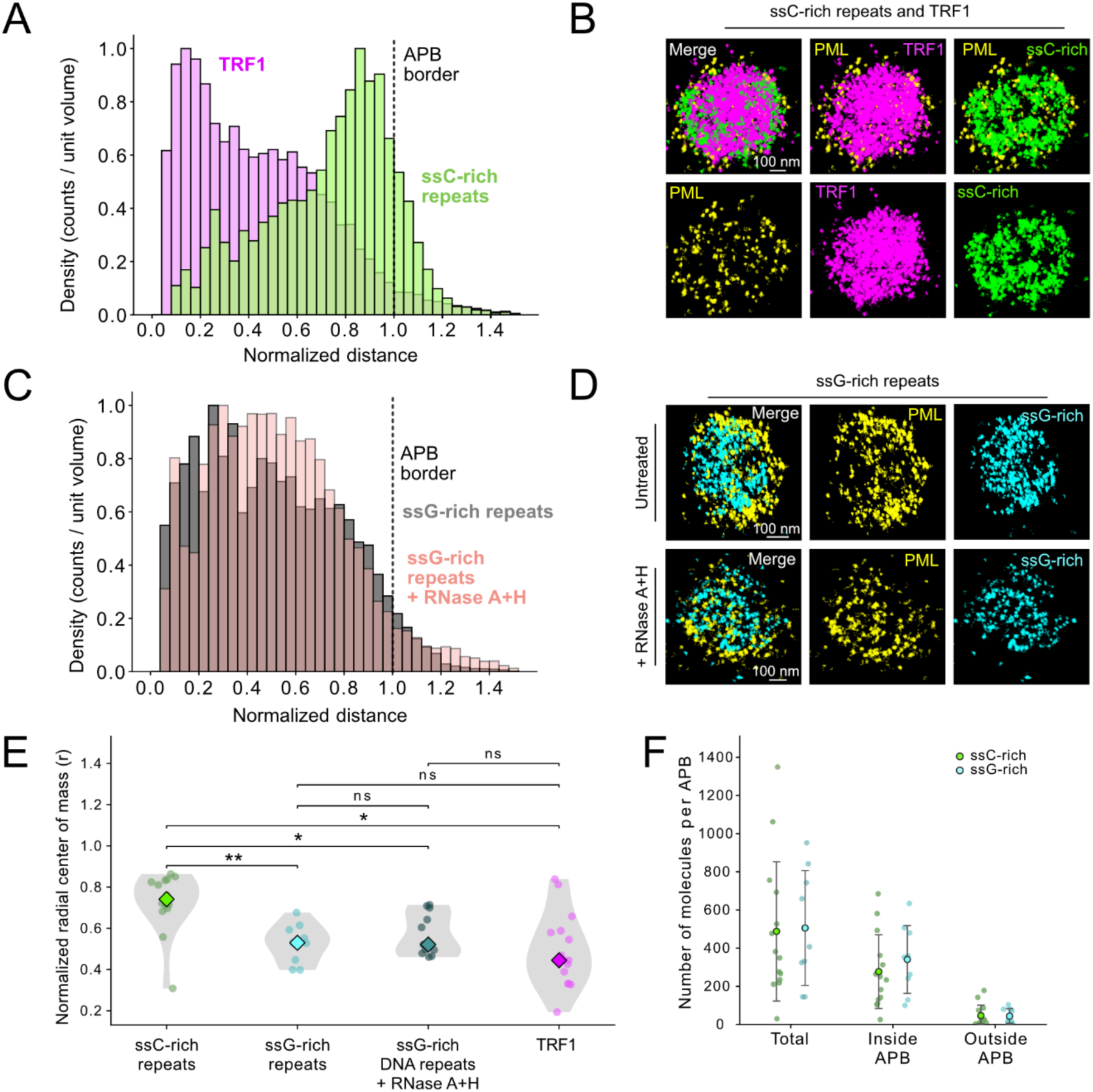
Radial organization of single-stranded telomeric repeats within APBs. (**A**) Volume-normalized radial density distributions of TRF1 (magenta, n = 14) and ssC-rich repeats (green, n = 14) relative to the APB center. Each distribution is independently normalized to 1 at its highest abundance peak. The dashed line indicates the PML shell border (r = 1). Histograms are normalized to counts per unit volume. (**B**) Representative 3D MINFLUX visualization of 3-target APB acquisition showing TRF1 (magenta), ssC-rich repeats (green), and PML (yellow). See also Supplementary Video 5-8. Scale bar, 100 nm. (**C**) Volume-normalized radial density distributions of ssG-rich repeats under untreated conditions (gray, n = 10) and following RNase A+H treatment (pink, n = 11). Each distribution is independently normalized to 1 at its highest abundance peak. The dashed line indicates the PML shell border (r = 1). (**D**) Representative 3D MINFLUX visualization of individual APBs showing PML (yellow) and ssG-rich repeats (cyan) under untreated conditions (top) and following RNase A+H treatment (bottom). Scale bar, 100 nm. (**E**) Comparison of the normalized radial center of mass of the localization distribution for ssC-rich repeats (n = 14), ssG-rich repeats (n = 10), ssG-rich repeats after RNase A+H treatment (n = 11), and TRF1 (n = 15). Violin plots show the distribution of individual APB structures (points); diamonds indicate the median. Group differences were first assessed using a Kruskal–Wallis test, followed by pairwise Mann–Whitney U tests with Bonferroni correction (* p < 0.05, ** p < 0.01, ns = not significant). (**F**) Quantification of ssC-rich (green) and ssG-rich (cyan) molecule counts per APB, categorized as total, inside APB, and outside APB. Point plots show mean ± SD; individual points represent single structures (ssC-rich n = 14, ssG-rich n = 10).

### ssG-rich repeats in APBs are predominantly DNA rather than unbound TERRA RNA

To directly assess the contribution of RNA to the ssG-rich repeat signal within APBs, we performed RNase A and H treatment. Successful RNase digestion was confirmed by a decrease in spot count and intensity of ssG-rich repeats, as reported previously (**Supplementary Fig. 5C, D**) ^8^. To quantify the unbound RNA fraction within APBs specifically, we compared confocal ssG-rich signal intensity inside APBs before and after RNase treatment using ALT-FISH. The reduction of 14% (p = 0.058) indicates that most of the ssG-rich signal within APBs is RNase-resistant (**Supplementary Fig. 5E**). Consistent with this, MINFLUX imaging after RNase treatment did not yield a detectable reduction in the number of localizations inside APBs, nor did it alter their radial distribution (p = 0.30; **Fig. 4E**). Together, these data demonstrate that the ssG-rich signal within APBs represents predominantly DNA rather than unbound TERRA RNA.

### ssG-rich DNA repeats frequently colocalize with POT1 assemblies

To further characterize the single-stranded DNA component inside APBs, we performed two-target MINFLUX imaging of GFP-POT1 and ssG-rich repeats (**Fig. 5A**). POT1 molecules were more abundant than ssG-rich repeat molecules, consistent with POT1’s dual role as a shelterin component and as a binder of single-stranded G-rich overhangs. NN analysis from ssG-rich repeats to the closest POT1 molecule revealed a peak distance of 16.5 nm (**Supplementary Fig. 6A**), indicating that a substantial fraction of ssG-rich repeats localizes in close proximity to POT1. The reciprocal analysis from POT1 to ssG-rich repeats yielded a larger peak distance (38.5 nm), reflecting that many POT1 molecules are not directly associated with ssG-rich DNA (**Supplementary Fig. 6B**). This asymmetry is consistent with the presence of POT1 within the shelterin complex independently of its ssDNA-binding function. Further analysis of ssG-rich to POT1 NN distances revealed a multimodal distribution that could be approximated by a three-component Gaussian model (**Fig. 5B**). The first component corresponds to ssG-rich repeats in close proximity to POT1 and is consistent with direct binding to single-stranded DNA. Additional components at larger distances may reflect distinct populations, such as TERRA RNA or non-specific background signal. Together, these data establish that a substantial fraction of ssG-rich repeats within APBs are bound by POT1, identifying them as single-stranded G-rich DNA substrates.

**Fig. 5.**
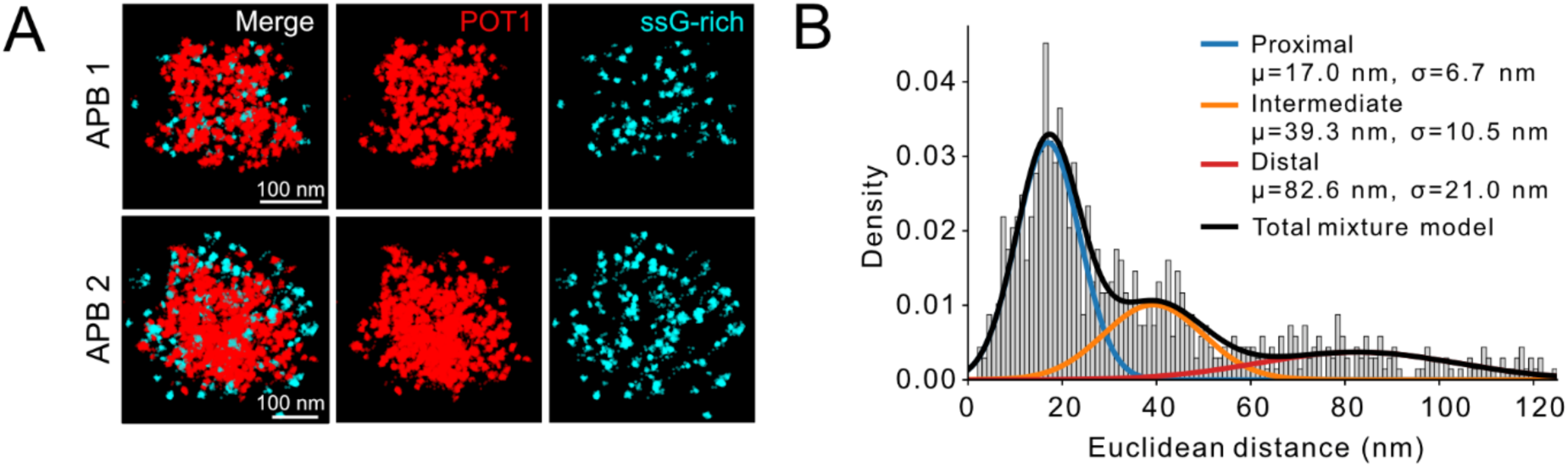
Single-stranded telomeric repeat localization in APBs. (**A**) Representative 3D MINFLUX visualization (shown as 2D projection) of two individual APBs showing the spatial organization of ssG-rich repeats (cyan, detected with (CCCTAA)₅ probe) relative to POT1 (red, detected with anti-GFP single-domain antibody). Scale bar, 100 nm. (**B**) First nearest-neighbor (NN) Euclidean distance distribution from ssG-rich repeats to POT1 (gray histogram, n = 4). A three-component Gaussian mixture model (black) was fitted to the data, identifying three subpopulations: a proximal population consistent with ssDNA-bound POT1 (blue), an intermediate population (orange), and a distal background population (red). Fitted means (µ) and standard deviations (σ) are indicated in the panel.

## Discussion

We resolved APB architecture at ∼3 nm isotropic precision by combining MINFLUX imaging with multiplexed exchange DNA-PAINT. This analysis adds a spatial dimension to mechanistic models of ALT, a heterogeneous process that can proceed through multiple parallel pathways ^34–36^. Our study reveals that ssC-rich repeats are enriched within the PML shell and that ssG-rich repeats with TRF1 are concentrated at the core (**Fig. 6A,B**).

**Fig. 6.**
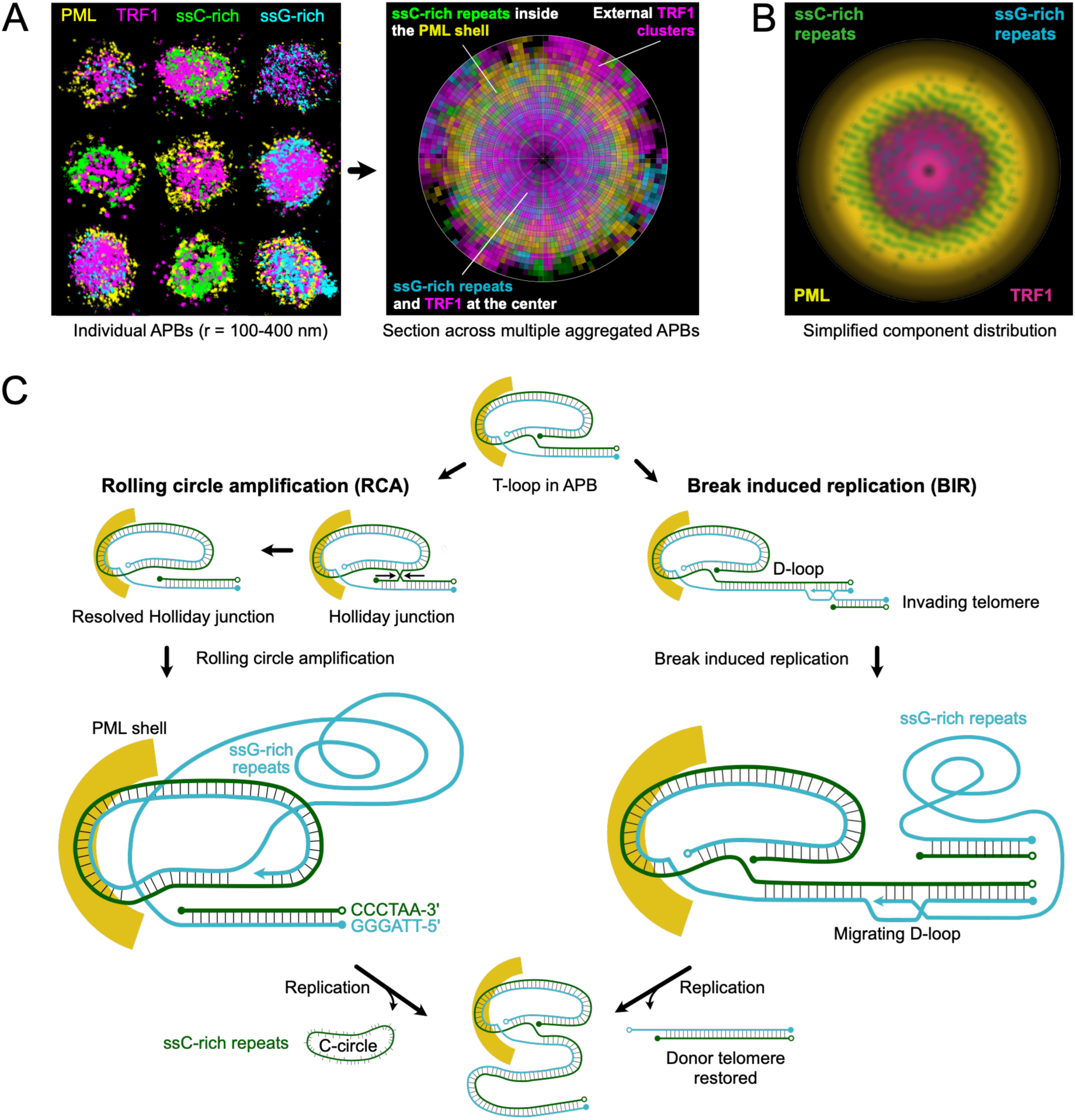
MINFLUX-derived APB organization and proposed routes for ALT telomere elongation. (A) Integrated model of APB nanoscale organization derived from 3-target MINFLUX imaging of individual APBs, shown as a 2D projection on the left and normalized to the same PML shell radius. The cross-section of an APB on the right was generated by aggregating 3-target images (5 with ssC-rich repeat staining and 6 with ssG-rich repeat staining) into a 4-target APB visualization. TRF1 was enriched toward the APB center, with clusters indicated outside the shell; ssG-rich repeats were broadly distributed across the interior; and ssC-rich repeats were enriched at the inner PML shell surface. Color coding: PML (yellow), TRF1 (magenta), ssG-rich repeats (cyan), and ssC-rich repeats (green). (B) Coarse-grained illustration of the distinct spatial compartmentalization of each component within the APB. (**C**) Two proposed routes for ALT-mediated telomere elongation are shown from a common t-loop intermediate. In the RCA branch, Holliday junction formation and resolution generate a circular template. Rolling-circle copying uses a C-circle template to extend the recipient telomere while displacing an ssG-rich strand. In the BIR branch, invasion of a homologous donor duplex forms a migrating D-loop that supports long-tract synthesis, restoring the donor and extending the recipient telomere. Nascent synthesis tracts in human ALT cells range from about 5 to 70 kb, with a median of about 20 kb ^37^. ssC-rich DNA is green, ssG-rich DNA is cyan, and PML is yellow.

It imposes structural constraints on two telomere-elongation pathways proposed for the ALT mechanism: rolling-circle amplification (RCA) on an extrachromosomal telomeric circle and break-induced replication (BIR) using a homologous duplex donor. The two routes differ in the mode of synthesis rather than in the template alone, since an extrachromosomal telomeric circle can also serve as the invaded donor of a conservative BIR reaction ^38^. Evidence for BIR comes from conservatively replicated telomeric DNA detected at a subset of telomeres in ALT-positive cells that depends on the polymerase delta subunits PolD3 and PolD4, two subunits of human DNA polymerase delta that are essential for BIR ^39^. Furthermore, a POLD3-dependent long-tract synthesis in an acute telomere-break system was shown ^37^. Although direct evidence for RCA in human cells is lacking, C-circles are a hallmark of ALT ^19^, and circle-templated copying stabilizes telomeres in yeast ^40^, indicating that RCA-competent templates are present. RCA and BIR are therefore best considered parallel, potentially coexisting contributions. This is depicted in our model, with both branches originating from a common t-loop-derived intermediate (**Fig. 6C**).

The APB architecture resolved here provides the structural context for the ALT reaction to proceed. Rather than the homogeneous liquid phase expected for a phase-separated condensate ^41^, PML in APBs forms a partially open shell with a Gaussian-shaped radial distribution and a lower density than canonical PML-NBs. Within the PML shell, ssC-rich repeats, ssG-rich repeats, and TRF1-marked telomeric chromatin occupy distinct radial positions across individual APBs (**Fig. 4A-E**, **Fig. 6A, B**), a pattern that would not be maintained by free mixing of components. The PML scaffold could direct component accumulation at its inner and outer surfaces, facilitate exchange of circle templates proposed for RCA, and bring telomeres into proximity to enable intertelomeric BIR.

TRF1-marked double-stranded telomeric chromatin occupies the shell interior, and one to three additional TRF1 clusters lie outside it, with exceptions in a small fraction of APBs. This arrangement brings several telomeric domains into proximity, in line with previous studies ^33,42^. Directional homology search and synapsis between ALT telomeres have been visualized in human ALT cells after induction of telomeric double-strand breaks ^43^, and telomere-derived sequences can be copied from one chromosome end to another ^5^. Thus, the clustered arrangement of TRF1 resolved by MINFLUX indicates donor proximity that a BIR-type reaction could exploit ^37^.

The ssC-rich repeats accumulated at the inner PML shell surface as a discrete pool, with abundance that scaled only weakly with shell size (ρ = 0.32) compared with TRF1 and ssG-rich repeats, suggesting an interaction with the PML scaffold. It could involve direct binding of ssDNA by PML ^44^ or indirect retention through shell-associated RPA ^45^, which preferentially binds ssC-rich DNA ^46^. Within APBs, POT1 coating or G-quadruplex formation could limit access to ssG-rich repeats and shift the accessible binding equilibrium toward ssC-rich DNA. Consistent with an RCA origin, ssC-rich telomeric DNA in ALT cells is largely circular, whereas ssG-rich repeats are predominantly linear ^47^. In the RCA model, branch migration and resolution of t-loop or i-loop junctions could release telomeric circles ^48,49^. Separate lagging-strand processing routes generate C-rich ssDNA and C-circles in ALT cells, whereas damage-induced i-loop resolution can also release telomeric circles ^10,50,51^. Copying a C-circle displaces an ssG-rich repeat strand, providing a direct mechanistic link between the two species observed in APBs (**Fig. 6C)**. Under a BIR extension model, C-circles would be generated separately through lagging-strand processing or loop excision, or as a product of the RAD52-independent BIR subpathway ^34^.

The broadly distributed ssG-rich repeats provide a second distinguishing feature between the two pathways. We have found that ssG-rich repeats are distributed across the same interior region as TRF1-marked double-stranded telomeric chromatin, were predominantly DNA rather than TERRA RNA, and frequently associated with POT1. In RCA, a G-rich single strand is displaced as the C-circle is copied, so the generation of ssG-rich DNA follows directly from the reaction (**Fig. 6C**). A BIR explanation would invoke a migrating D-loop that displaces the newly synthesized strand (**Fig. 6C**). This geometry is established for non-telomeric break-induced replication in budding yeast, where a Pif1 and polymerase delta driven migrating D-loop displaces the newly synthesized strand and yields conservative inheritance of the new DNA ^52,53^. Whether the same geometry operates on telomeric repeat arrays in human APBs, which additionally depend on PML, a protein without a yeast counterpart, remains to be demonstrated. Distinguishing between the BIR and RCA route experimentally will require resolving nascent DNA synthesis within the APB. Labeling nascent DNA and resolving it against the ssC-rich pool and the PML shell would test whether C-circles are made at the inner shell or generated elsewhere and only sequestered there. Directly linking nascent telomeric DNA either to a second chromosomal telomere or to a circular extrachromosomal template would distinguish the intertelomeric BIR and circle-templated RCA routes drawn in **Fig. 6C**.

By resolving APB architecture at the molecular scale, multiplexed 3D MINFLUX provides a strategy for defining the nanoscale organization of telomeres in cells that employ ALT. The same approach could be extended to map the molecular architecture of the telomerase-maintained telosome at individual chromosome ends. More broadly, the combination of 3D MINFLUX with exchange DNA-PAINT establishes a general framework for resolving the molecular architecture of nuclear compartments in situ and constraining the mechanisms that govern their organization and function.

## Acknowledgements

This project was supported by the Cooperation Program in Cancer Research of the German Cancer Research Center (DKFZ) and Israel’s Ministry of Science, Technology and Space (MOST) via grant Ca215 to KR. The MINFLUX nanoscope is part of the CellNetworks Core Technology Platform (CCTP) of Heidelberg University and was financed by the European Union’s fund for regional development (EFRE) -innovation and energy change as part of the reaction to the COVID-19 pandemic (REACT-EU) with upgrades and extensions funded by the Federal Ministry of Research, Technology, and Space (BMFTR) and the Ministry of Science Baden-Württemberg (MWK) within the framework of the Excellence Strategy of the Federal and State Governments of Germany. Data storage at SDS@hd was funded by the MWK and the German Research Foundation (DFG) through grant INST 35/1503-1 FUGG.

## Author contributions

Acquisition of data: EK, CK, CaK, DB. Analysis and interpretation of data: EK, ML, CK, KR. Design and conceptualization: EK, KR. Writing of the original draft: EK, KR. Reviewing & editing of manuscript: all authors. Supervision: KR.

## Methods

### Cell culture

All cell lines were cultured at 37 °C in 5% CO2 and split every 2-4 days using 0.05% trypsin/EDTA (Gibco, Thermo Fisher Scientific, Waltham, MA, USA) for detachment. U2OS cells were cultured in DMEM containing 1 g/L glucose (Gibco) supplemented with 10% doxycycline-free fetal bovine serum (PAN-Biotech, Aidenbach, Germany), 1x penicillin/streptomycin (PAN-Biotech), and 2 mM stable glutamine (PAN-Biotech). U2OS WT cells were obtained from the German Collection of Microorganisms and Cell Culture (DSMZ, Germany). The identity of all cell lines was confirmed by SNP-profiling-based cell line authentication (Multiplexion, Germany). All cells tested negative for mycoplasma (VenorGeM Advance, Minerva Biolabs, Berlin, Germany).

### Cell line generation

U2OS HaloTag-PML TRF1-eGFP cells were created by first establishing stable cell lines with doxycycline-inducible TRF1-eGFP expression using the Tet-On 3G system (Clontech, Takara Bio, Kusatsu, Japan). U2OS Tet-On 3G cells were generated by stable transfection of the geneticin-resistant pCMV-TET3G plasmid, followed by selection with 1 mg/mL G-418 (Thermo Fisher Scientific) to obtain plasmid-carrying cells. Cells were then stably transfected with the puromycin-resistant pTRE3G-TRF1-eGFP plasmid. This plasmid was generated by cloning TRF1-eGFP from a TRF1-eGFP plasmid created using a pEGFP-N1 plasmid (Clontech) with TRF1 inserted by digestion with SacI and NheI. The TRF1-eGFP plasmid was inserted into a pTRE3G plasmid through digestion with SalI and PstI. U2OS WT cells were transfected with the pTRE3G-TRF1-eGFP plasmid using the Xfect transfection reagent (Clontech). After 48 h, the cells were expanded, and after another 48 h, G-418 (1 mg/mL) and puromycin (0.5 µg/mL, Sigma-Aldrich, Merck KGaA, Darmstadt, Germany) were added. After approximately 14 days, when G-418-resistant colonies appeared, single colonies were picked and tested for proper induction of TRF1-eGFP using doxycycline (100 ng/mL, Sigma-Aldrich).

### Endogenous tagging

After validation, CRISPR/Cas9 was used to endogenously tag the N-terminus of PML with a HaloTag (Promega, Madison, WI, USA). A 3xHA-HaloTag insert was obtained as a gBlock (IDT, Coralville, IA, USA) containing PML-specific 90-bp flanking genomic DNA sequences at each site. Additionally, a mutation in the PAM sequence was introduced to prevent re-cutting by pSpCas9 after the initial break repair. The gBlocks were cloned into a KpnI-XbaI-digested pUC19 vector (Addgene, Watertown, MA, USA; #50005) by Gibson assembly, yielding a pUC19-3xHA-HaloTag-TEV plasmid. This plasmid was nucleofected together with Alt-R S.p.Cas9 Nuclease V3 (IDT) and an Alt-R CRISPR-Cas9 crRNA (IDT) targeting the correct integration site. The crRNA (5’-UCGGGCGGGUGCAGGCUCCAGUUUUAGAGCUAUGCU-3’) was designed to overlap with the PML start codon using CRISPOR and BLAST.

First, cells were treated for 48 h with Lipofectamine RNAiMAX (Invitrogen, Thermo Fisher Scientific) and 5 nM p53 siRNA pool (siTOOLS Biotech, Planegg, Germany) to support cell cycling, and with 1 mM M3814 (MedChemExpress, Monmouth Junction, NJ, USA) to inhibit non-homologous end joining (NHEJ). Then, nucleofection was performed on U2OS stable cell lines with inducible TRF1-GFP using a 4D-Nucleofector (Lonza, Basel, Switzerland) with the SE cell line 4D nucleofector X kit (Lonza). The CM-104 program was used.

For nucleofection, an RNP complex consisting of the gRNA (30 µM) and Alt-R Cas9 enzyme (IDT), together with an Alt-R electroporation enhancer, was used with 5 µg of plasmid DNA. After 12 h, the medium was refreshed, and cells were expanded for FACS sorting. To generate monoclonal endogenously tagged cell lines, cells were incubated with the Janelia Fluor HaloTag 646 ligand (Promega) and then single-cell sorted into a 96-well plate based on fluorescence using a BD FACSAria Fusion flow cytometer. After 7-9 days, all wells were checked to select those with single colonies, which were then grown out and tested by PCR outside the plasmid homology arms to identify clones with an endogenous HaloTag (FWD: 5’-CCTACCTCTCCCGCTTTACC-3’ and REV 5’-CACCTCGGATCTGGGAAGTT-3’). PCR bands were sequenced to confirm that no mutations were introduced at the insertion site. Expression of the tagged protein was confirmed by western blot, and immunofluorescence PML Ab staining was combined with Halo-JF646 incubation to confirm cellular localization.

### ALT-FISH with RNase treatment

U2OS cells with endogenous HaloTag-PML and inducible TRF1-GFP were plated on 12-mm coverslips, and TRF1-GFP expression was induced 16 h before fixation by adding doxycycline. Cells were incubated for 20 min with 200 nM HaloTag Janelia Fluor 549 ligand (Promega) and then washed three times for 5 min with pre-warmed medium at 37°C. Cells were then washed twice with PBS and fixed in 70% ethanol for 20 min at room temperature. After two washes with wash buffer (0.1 M Tris-HCl pH 8.0, 0.15 M NaCl, 0.05% Tween-20), cells were incubated for 30 min at 37°C with either 50 µg/mL RNase A (Thermo Fisher Scientific) and 10 U/mL RNase H (New England Biolabs, Ipswich, MA, USA) in PBS, or with 10 mM vanadyl ribonucleoside complex (VRC) as an RNase inhibitor control. After one wash with wash buffer, cells were incubated with 5 nM ALT-FISH ssC-rich probe (5′-(CCCTAA)₅-3′-Biotin, labeled with Atto633) for 20 min at 37°C, followed by two washes with 2× SSC. Cells were then incubated for 15 min with 5 µM DAPI in PBS, washed three times in PBS, briefly rinsed sequentially with water, 70% ethanol, and 100% ethanol, air-dried, and mounted in ProLong Diamond Antifade Mountant (Invitrogen, Thermo Fisher Scientific). Coverslips were stored overnight in the dark before imaging.

### Spinning disk confocal microscopy

Confocal fluorescence imaging was performed on an Andor Dragonfly 505 spinning-disk confocal microscope (Andor Technology, Belfast, UK) mounted on a Nikon Ti2-E inverted stand (Nikon, Tokyo, Japan) using a 100× silicone-immersion objective (CFI SR HP Apochromat Lambda S 100×, Nikon). Excitation was provided by laser lines at 405 nm, 488 nm, 561 nm, and 637 nm, with a quint-band dichroic mirror (405/488/561/640/750 nm) and emission filters at 445/46, 521/38, 571/78, and 685/47 nm. Fluorescence was detected with an iXon Ultra 888 EM-CCD camera (Andor Technology), and images were acquired at 16-bit depth with a field size of 1024 × 1024 pixels (pixel size: 120.5 nm).

### MINFLUX sample preparation

Approximately 2 × 10^5^ U2OS cells were grown in uncoated 0.8 mm μ-Slide Luer (Ibidi, Gräfelfing, Germany) channel slides to 70-80% confluence. For U2OS HaloTag-PML TRF1-GFP cells, 1 µg/ml doxycycline hyclate (Sigma-Aldrich) was added at plating to induce TRF1-GFP expression. The next day, cells were incubated for 45 min with 200 nM Janelia Fluor 646 HaloTag ligand (Promega) and then washed three times in PBS. Cells were fixed for 10 min in 4% PFA (Sigma-Aldrich), washed three times with PBS, permeabilized for 5 min with 0.5% Triton X-100 (Merck) in PBS, and washed three times with PBS. Subsequently, probe hybridization was performed in hybridization solution (2x SSC, de-ionized formamide, 50 nM probe) and incubated in a humidified chamber for 2 h at 37 °C, using either a C-rich (Docking strand 4-AAAA-(TTAGGG)_5_) or G-rich (Docking strand 2-AAAA-(CCCTAA)_5_) telomeric probe consisting of telomeric repeats, a flexible linker, and a docking strand (Massive Photonics, Munich, Germany). After hybridization, the sample was washed first for 5 min with 2X SSC/50% formamide, followed by two 5 min washes with 0.1% Tween in 2x SSC buffer, and incubated for 1 h in PBS containing 10% goat serum (blocking buffer) (Cell Signaling Technology, Danves, USA). Then, the sample was incubated with the primary antibody for 1 h at RT (**Supplementary Table 1**). After three washes with 0.002% NP-40/PBS, samples were washed once with washing buffer (Massive Photonics), followed by a 1 h incubation with DNA-PAINT sdAbs (**Supplementary Table 1**). The sample was then washed three times in washing buffer and stored in PBS at 4 °C in the dark until imaging. For POT1 transfection, U2OS HaloTag-PML TRF1-GFP cells were plated in an uncoated 0.8 mm μ-Slide Luer (Ibidi) channel slide in the absence of doxycycline. After attachment, cells were transfected with 1 µg GFP-POT1 plasmid in Opti-MEM (Gibco) using Xtreme Gene HP9 (Sigma-Aldrich), following the manufacturer’s recommendations. After 24 h, cells were fixed and stained as described before.

### Gold bead preparation

On the day of MINFLUX measurements, one-target samples were incubated for 5 min with 150 nm gold beads (A11-150-CIT-DIH-1, Nanopartz, Loveland, US), then washed twice in 1x PBS (D8537, Sigma-Aldrich). Samples were incubated with 0.1% poly-L-lysine (P8920, Sigma-Aldrich) for 3 min, then washed again in 1x PBS before the imaging buffer containing the respective imager was added (**Supplementary Table 2**; Massive Photonics). Three-target samples were prepared by incubating for 5 min with a 1:2 mixture of 150 nm and 200 nm gold beads (A11-200-CIT-DIH-1, Nanopartz). They were then washed three times in 1x PBS, incubated with poly-L-lysine for 5 min, and washed four times in 1x PBS. Before mounting on the stage, imaging buffer (Massive Photonics) was added. DNA-PAINT imagers (Im1, Im2, Im3, and Im4) conjugated to a Cy3B fluorophore and dissolved in TE buffer (10 mM Tris, 1 mM EDTA, pH 8) were purchased from Massive Photonics.

### MINFLUX microscopy

A MINFLUX microscope (Abberior Instruments, Göttingen, Germany) was used for all MINFLUX single-molecule localization experiments. The MINFLUX microscope was built on a motorized inverted Olympus IX83 microscope (Evident Europe, Hamburg, Germany) equipped with a 100x 1.45x UPLanXAPO Olympus oil objective, four laser lines: 405 nm, 488 nm, 561 nm, and 640 nm, and a COOL LED pE-4000 for epifluorescence illumination. The MINFLUX microscope was controlled by the Imspector software (versions 16.3.15645-m2205 and 16.321317-m2410). More details on the hardware components of the MINFLUX setup were described previously ^54^. MINFLUX measurements were performed with high-precision sample and laser drift correction. Sample drift was corrected by active stabilization using a 980 nm IR laser and an XYZ piezo stage (Piezoconcept, Bron, France). Camera images of the backscattered light from gold beads applied to the sample beforehand were used in a control loop to correct the piezo stage position with sub-nanometer precision in x, y, and z. For laser drift correction, the beam line monitoring system was used on at least four gold beads during MINFLUX measurements. Three-dimensional MINFLUX localization measurements were performed with the MINFLUX sequence provided by the manufacturer (**Supplementary Table 3**). For all measurements, the 561 laser line was used at a power of 27 µW to 38 µW (measured at the periscope), and detection was performed with an APD with a spectral window of 580 nm to 630 nm. The pinhole size was set to 0.83 airy units. Before each MINFLUX acquisition, confocal images were acquired for each channel, using a pixel size of 60 for an overview image of the cell, followed by an acquisition of the structure of interest with a pixel size of 10.

### Automated Exchange-PAINT MINFLUX measurements

For multitarget imaging by Exchange-PAINT, a commercial automated perfusion system (Aria M2, Fluigent, Le Kremlin Bicetre, France) was used in combination with a microfluidics channel µ-Slide Luer with a #1.5 high-precision glass bottom (80197, Ibidi). The Aria system was controlled by the Aria software (v2.3.1.0). Solutions containing the respective imagers (**Supplementary Table 2**) were prepared in 15 mL centrifuge tubes and mounted on the Aria. Imaging buffer (Massive Photonics) was used as the wash buffer. The same protocol for automatically exchanging the solution in the sample during the MINFLUX measurements was used for all three-target measurements (**Supplementary Table 4**). Additionally, the same protocol was used for all 2-target experiments (**Supplementary Table 5**). All solutions were exchanged at a rate of 30 µL/min to maintain active sample stabilization and beamline monitoring throughout the MINFLUX measurement. MINFLUX measurements were started once the imager solution was completely filled into the channel slide, before the “2 hours waiting” step. The MINFLUX measurement continued throughout the protocol to ensure that no localizations were detected before the sample floated into the next imager solution.

### MINFLUX image reconstruction and visualization

Visualizations of MINFLUX images were generated using Paraview (V5.12.1) ^55^. The .npy files were exported from Abberior Imspector software (V16.3.21317-m2410) and converted to .pmx files using PyMINFLUX (V0.0.6). For visualization of localizations in Paraview, the representation was set to point gaussian, with a gaussian radius of 4 nm. The shader preset was plain circle. Visualizations were set to an opacity of 0.4, specular was set to 0, and the scaling mode was set to all approximate.

### MINFLUX data filtering and analysis

All MINFLUX data were exported as .npy files, and data analysis was performed using Python (v.3.11.10) with the following packages: numpy (v1.24.3), matplotlib-base (v3.10.7), scikit-learn (v1.7.2), seaborn (v0.13.2), pandas (v2.2.3), and scipy (v1.15.3). For the analysis, only valid x, y, and z coordinates from the ninth iteration of a MINFLUX measurement, referred to as localizations, were considered. Individual traces were analyzed to identify multiple localization distributions or tail artifacts. For each trace, the localization standard deviation was computed. If the trace’s standard deviation exceeded three times the combined localization precision, defined as the square root of the sum of the squared localization precisions in x, y, and z, the trace was subjected to a DBSCAN clustering algorithm with a minimum point count of three and a search radius equal to the combined localization precision. Localizations labeled as noise by DBSCAN were excluded from the trace. In cases where multiple distinct localization populations were identified within the same trace, new trace identifiers were assigned to the additional populations. This approach effectively mitigates tail artifacts and enables the detection of multiple localization populations within individual traces. A scaling factor of 0.7 was applied to the z-coordinates to compensate for the apparent axial elongation due to refractive index mismatch between the oil immersion lens and the sample medium ^56^.

To automatically split MINFLUX measurements when two or more exchange DNA-PAINT targets were imaged in a single measurement file, we developed an analysis based on valid event rates. For each measurement file, the second derivative of the number of valid localizations per minute was calculated. A lower and higher bound were defined as lower bound = 1^st^ Percentile – K *(99^th^ Percentile – 1^st^ percentile) and higher bound = 99^th^ Percentile + K *(99^th^ Percentile – 1^st^ percentile), where K is a tuning factor varying between 1 and 2. The time when the signal crossed the higher bound was defined as the buffer end time, and whenever the signal crossed the lower bound, it was defined as the buffer beginning time. This method successfully split the experiment according to the imager buffer used for most datasets. When automatic splitting was not possible due to heterogeneous event rates between different imager buffers, the time thresholds were defined manually. The K threshold was set based on the number of valid localizations per minute, and each dataset was then saved as a .npy file. Raw valid localization data was aggregated by a photon count of 500 in Imspector and exported as a .npy file. These files underwent the same data analysis procedure as described above.

### Analysis of PML spherical shell

For the analysis of the PML shell, the target structure was isolated by applying the DBSCAN clustering algorithm from scikit-learn (version 1.7.2) to the centroid positions of traces, computed as the mean of all localizations associated with each trace. DBSCAN parameters were optimized per acquisition: the minimum number of points was set between 50 and 200, while the search radius (epsilon) ranged from 120 to 300 nm to accommodate structural heterogeneity. Traces identified as noise by DBSCAN, i.e., those not belonging to the target structure, were excluded from subsequent analyses. To model the PML structure, a least-squares sphere fit was performed on the remaining trace centroids to estimate the structure’s center coordinates and radius by minimizing residuals. PML shell thickness was determined by calculating the Euclidean distances from each trace centroid to the fitted sphere center, which were then binned into 5 nm intervals. Local minima (valleys) in the resulting histogram were identified using SciPy’s ‘find_peaks’ function on the inverted histogram data, applying a peak density threshold of 0.001. The first and last valleys were designated as the inner and outer radii of the PML shell, respectively. Next, to compare distributions of molecules across the APB while accounting for size differences, molecule counts were normalized to the corresponding shell volume, assuming a spherical shell geometry. Each shell was defined by an inner radius r (given by the fitted sphere radius) and an outer radius r + t, where t denotes the shell thickness. The shell volume was calculated as

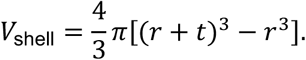

Volume-normalized densities were defined as the number of molecules per unit shell volume (molecules·nm^-3^). A normalized Gaussian mixture model was used to describe the distribution of PML localizations across the shell structure.

### Protein nearest-neighbor and clustering analysis

For nearest-neighbor (NN) distance analysis, the 5 nearest neighbors were computed for each molecule using the Euclidean metric. Distances were also grouped by relative position to the APB: inside the APB, within the shell, and outside the APB structure. The haversine distance, defined as the shortest great-circle arc length between molecular positions on a spherical surface with an assumed radius, was also used. However, because the molecular shell has finite, non-negligible thickness and a spatial distribution within this volume, the computed haversine distances were significantly influenced by the assumption of a zero-thickness (idealized spherical surface). Therefore, given the spatial distribution of molecules within the shell volume, Euclidean distance was the most suitable metric for NN analysis. Distributions were visualized using kernel density estimation (KDE) and normalized histograms. Statistical differences between conditions were assessed using the two-sample Kolmogorov–Smirnov (KS) test (two-sided), which compares the full empirical cumulative distribution functions. The p-values were adjusted using Bonferroni correction for multiple testing across NN orders. To identify possible clustering of TRF1 and single-stranded telomeric repeat structures, DBSCAN was applied to the centroid positions of molecules lying outside the APB shell. The minimum number of molecules to be considered a cluster was 5, and the search radius was 80 nm. For each identified cluster, the center was calculated as the arithmetic mean of all three-dimensional localization coordinates belonging to that cluster.

### Angular and radial distribution analysis

The spatial distribution of telomere and TRF1 clusters was analyzed by converting cluster centroids from Cartesian to spherical coordinates and characterizing them with comparative angular analysis across samples. Azimuthal angles were rotated so that the cluster with the most telomere localizations was aligned to theta = 0. This was achieved by subtracting the azimuthal angle of the largest cluster centroid from all cluster angles. Polar plots were then generated to visualize radial distance as a function of azimuthal angle, enabling assessment of the angular distribution of telomere clusters relative to the PML-centered coordinate system.

Telomeric repeat localizations were quantified relative to the APB structure with normalized radial distance density plots. For each localization, the distance to the PML center and the corresponding PML radius were calculated. Then a normalized radial position was computed as 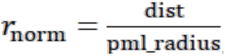. Localization distances were restricted to maximum r_norm_ = 1.5 to still be considered part of the APB structure. To account for the increasing volume at larger radii, acquisition-level radial profiles were computed as volume-normalized radial density. For each acquisition, *r*_norm_ values were binned with a bin width 0.04. Counts per bin were divided by the corresponding spherical shell volume in a unit sphere, 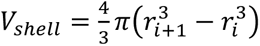, where *r_i_*_+1_ is the outer radius and *r_i_* the inner radius. This yielded a density profile (counts per unit volume) across radial bins for each acquisition.

For quantification, each acquisition-level density profile was summarized by its center of mass (CoM), computed as the density-weighted mean radial position:

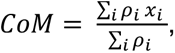

where *ρ_i_* is the volume-normalized density in bin *i* and *x_i_* is the corresponding bin center. The CoM was calculated independently for each acquisition, treating acquisitions as the unit of replication. Group differences in CoM between experimental conditions were assessed using non-parametric permutation tests on the difference in mean CoM values. Condition labels were randomly permuted across acquisitions (20,000 permutations), and the two-sided p-value was calculated as the proportion of permutations that produced an absolute mean difference at least as large as the observed difference (with +1 smoothing to avoid zero p-values).

### Confocal microscopy RNase treatment image analysis

Images were flat-field corrected and corrected for chromatic aberration before maximum intensity projection. Cell segmentation was performed using Cellpose3 ^57^. ALT-FISH spots were detected and counted using the RS-FISH plugin ^58^ in Fiji (v2.1.0; ^59^) with the following parameters: Sigma = 1.75, threshold = 0.0011, default RANSAC parameters, no background subtraction. Segmentation masks for TRF1-GFP and HaloTag-PML signals were generated using ilastik (v1.4.0; ^60^), and subsequent quantitative analysis was performed using a custom R pipeline (v2023.12.1, R v4.2.1; available at https://github.com/RippeLab/).

### TRF1-GFP dimer distance estimation

The average center-to-center distance between two GFP monomers fused to the C-termini of a TRF1 dimer was calculated using an AlphaFold2 structural model of the full-length TRF1-GFP dimer as the starting geometry. The distance between the C-termini of the MYB domains was 61.3 Å in this model. Each linker has a contour length of 3.8 nm (L ≈ 10 × 0.38 nm). The peptide persistence length in the random-coil regime is approximately 0.5–0.8 nm, yielding an RMS end-to-end extension per linker of approximately 2.0–2.5 nm. The offset from the GFP N-terminus to the GFP center of mass was approximated as 1.8 nm based on the barrel geometry of the GFP structure. A Monte Carlo simulation with two independent flexible linkers, random GFP orientations, and no steric constraints yielded a mean center-to-center GFP distance of 6.9–7.1 nm, an interquartile range of 5.5–8.7 nm, and a 5th–95th percentile range of 3.4–11.0 nm. The AlphaFold2 model predicted a center-to-center distance of 10.3 nm, which falls near the upper tail of the random-coil distribution.

## Data availability

The source images will be deposited at the BioImage Archive at https://www.ebi.ac.uk/bioimage-archive/S-BIAD3842. Custom analysis software is available on GitHub at https://github.com/RippeLab/MINFLUX-telomere-analysis.

