## Supplementary figures for "Single-stranded telomeric repeats segregate into spatial compartments within ALT-associated PML bodies"

### Content

**Supplementary tables**

Supplementary Table 1. List of antibodies used in this study.

Supplementary Table 2. DNA-PAINT imagers and the concentrations used.

Supplementary Table 3. MINFLUX sequence for 3D localization measurements.

Supplementary Table 4. ARIA software protocols for 3-target MINFLUX measurements.

Supplementary Table 5. ARIA software protocols for 2-target MINFLUX measurements.

**Supplementary figures**

Supplementary Figure 1. Generation and validation of the U2OS HaloTag-PML inducible TRF1-GFP cell line and MINFLUX quality parameters.

Supplementary Figure 2. Nearest-neighbor distance analysis of PML molecules in APBs compared to canonical PML nuclear bodies (PML-NBs).

Supplementary Figure 3. Photon count aggregation improves localization precision without affecting nearest-neighbor distances.

Supplementary Figure 4. Quantification of APB component scaling and TRF1 organization.

Supplementary Figure 5. Radial distribution of APB components across individual acquisitions and validation of RNase treatment efficacy.

Supplementary Figure 6. First nearest-neighbor (NN) distance analysis between POT1 and single-stranded (ss) G-rich repeats.

Supplementary Figure 7. Radial abundance profiles reveal spatial segregation of components within APBs.

**Supplementary videos**

Supplementary Video 1. Three-dimensional MINFLUX rendering of a representative APB with three targets.

Supplementary Video 2. PML channel of the APB shown in Supplementary Video 1.

Supplementary Video 3. TRF1 channel of the APB shown in Supplementary Video 1.

Supplementary Video 4. ssG-rich telomeric repeat channel of the APB shown in Supplementary Video 1.

Supplementary Video 5. Three-dimensional MINFLUX rendering of a second representative APB with three targets.

Supplementary Video 6. PML channel of the APB shown in Supplementary Video 5.

Supplementary Video 7. TRF1 channel of the APB shown in Supplementary Video 5.

Supplementary Video 8. ssC-rich telomeric repeat channel of the APB shown in Supplementary Video 5.

### Supplementary tables

**Supplementary Table 1. List of antibodies used in this study.**

| Antibody | Company | Number | Dilution |
|---|---|---|---|
| Mouse anti-PML | Abcam | ab96051 | 1:500 |
| sdAb Anti-Mouse IgG kappa light chain + Docking site 1 | Massive Photonics | | For PML: 1:200 |
| sdAb anti-GFP + Docking site 3 | Massive Photonics | | For TRF1-GFP: 1:500 |

**Supplementary Table 2. DNA-PAINT imagers and the concentrations used.**

| Target | Imager conjugated to Cy3B (1 µM stock) | Final concentration | Imager with no dye (10 µM stock) |
|---|---|---|---|
| PML | Imager 1 | 250 - 500 pM | |
| TRF1-GFP | Imager 3 | 125 pM | 8 nM |
| POT1-GFP | Imager 3 | 125 pM | |
| ssG-rich | Imager 4 | 500 pM | |
| ssC-rich probe | Imager 2 | 250 pM | 3.5 nM |

**Supplementary Table 3. MINFLUX sequence for 3D localization measurements.**

| Iteration | L (nm) | TCP parameter L(nm) | Photon threshold | Background threshold (Hz) | Pattern dwell time (ms) | Pattern repeat/s | Center frequency ratio upper limit | Laser power factor |
|---|---|---|---|---|---|---|---|---|
| 0 | 288 | Hexagon | 160 | 15000 | 1 | 1 | off | 1 |
| 1 | 1400 | z-line | 400 | 15000 | 1 | 1 | off | 1 |
| 2 | 288 | Square | 100 | 10000 | 1 | 5 | off | 1 |
| 3 | 288 | z-line2 | 50 | 10000 | 1 | 5 | off | 1 |
| 4 | 151 | Square | 67 | 10000 | 1 | 5 | 0.9 | 2 |
| 5 | 151 | z-line2 | 33 | 10000 | 1 | 5 | off | 2 |
| 6 | 76 | Square | 67 | 10000 | 1 | 5 | 0.8 | 4 |
| 7 | 76 | z-line2 | 33 | 10000 | 1 | 5 | off | 4 |
| 8 | 40 | Square | 100 | 10000 | 1 | 5 | off | 6 |
| 9 | 40 | z-line2 | 50 | 10000 | 1 | 5 | off | 6 |

**Supplementary Table 4. ARIA software protocols for 3-target MINFLUX measurements.**

| Steps | Action | Target |
|---|---|---|
| 1. | Fill 600 µL washing buffer | |
| 2. | Fill 300 µL solution with Imager 1 | PML |
| 3. | Wait for 2 h | |
| 4. | Fill 600 µL washing buffer | |
| 5. | Fill 300 µL solution with Imager 2 / 4 | ssG-rich, ssC-rich repeats |
| 6. | Wait for 2 h 30 min | |
| 7. | Fill 600 µL washing buffer | |
| 8. | Fill 300 µL solution with Imager 3 | TRF1-GFP |
| 9. | Wait minimum 3 h | |

**Supplementary Table 5. ARIA software protocols for 2-target MINFLUX measurements.**

| Steps | Action | Target |
|---|---|---|
| 1. | Fill 600 µL washing buffer | |
| 2. | Fill 300 µL solution with Imager 4 | ssG-rich repeats |
| 3. | Wait for 2 h | |
| 4. | Fill 600 µL washing buffer | |
| 5. | Fill 300 µL solution with Imager 1 | PML |
| 6. | Wait minimum 3 h | |

### Supplementary figures

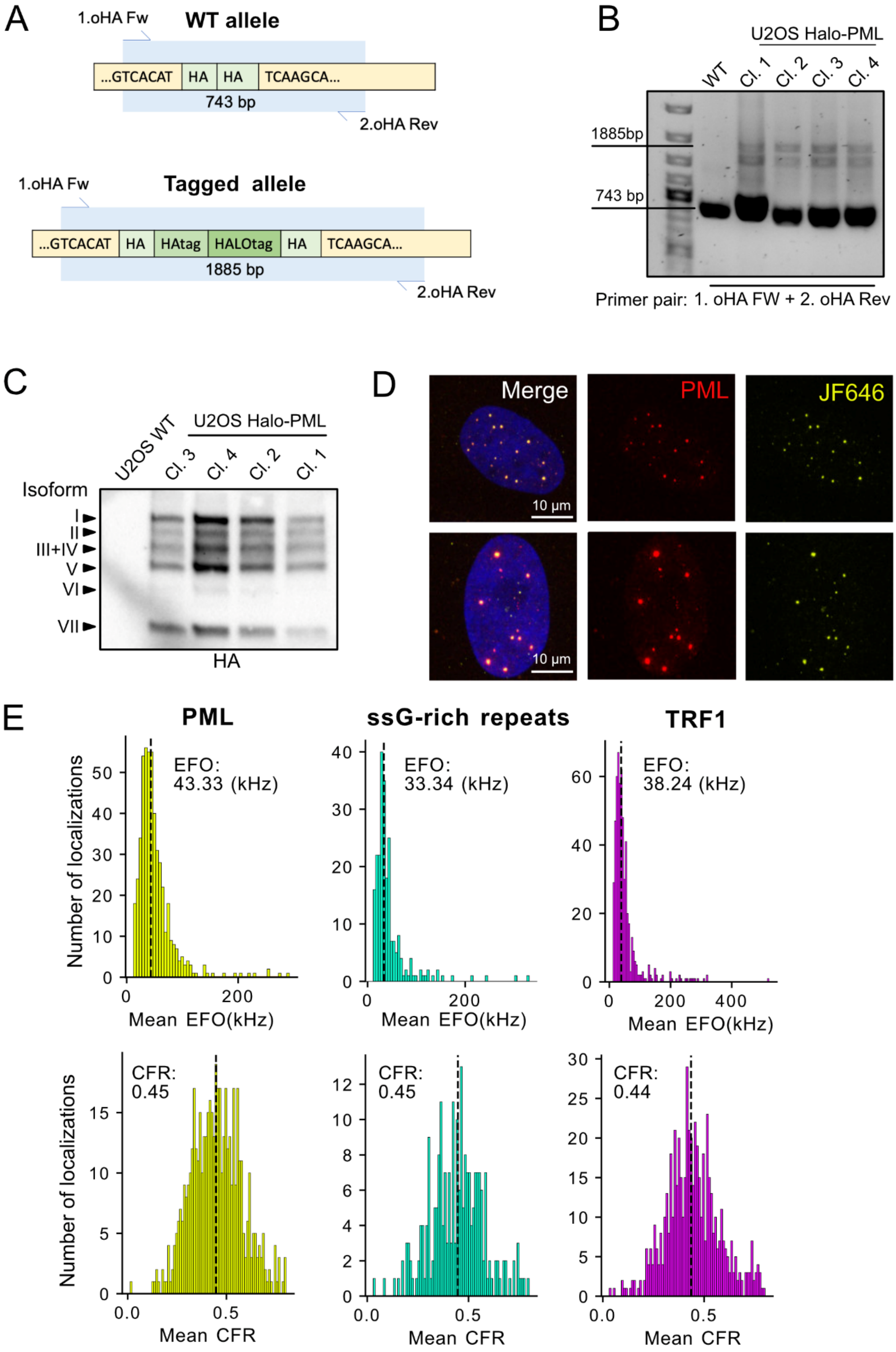

**Supplementary Figure 1. Generation and validation of the U2OS HaloTag-PML inducible TRF1-GFP cell line and MINFLUX quality parameters.** (**A**) Schematic overview of CRISPR/Cas9-mediated insertion of a HaloTag at the N-terminus of the endogenous PML locus, showing the primer pairs designed outside the homology arms (oHA) used for genomic validation. A wildtype (WT) allele yields a 743 bp PCR product, whereas a correctly targeted allele yields a 1885 bp product. (**B**) PCR validation of HaloTag-PML knock-in clones using primers outside the homology arms (oHA), confirming successful integration by the presence of the expected 1885 bp product. (**C**) Western blot using an anti-HA antibody confirming expression of HaloTag-PML across all seven PML isoforms in each validated U2OS clone. (**D**) Immunofluorescence confirming colocalization of anti-PML antibody signal with JF646-labeled HaloTag-PML. Scale bar, 10 µm. (**E**) MINFLUX quality parameters from a representative three-target exchange DNA-PAINT measurement of PML, ssG-rich repeats and TRF1-GFP, each detected with Cy3B-conjugated imager strands. Emission frequency offset (EFO) and center frequency ratio (CFR) values are shown per target. Mean EFO values were 43 kHz (PML), 33 kHz (ssG-rich repeats) and 38 kHz (TRF1-GFP), and mean CFR values were 0.44, 0.44 and 0.43, respectively.

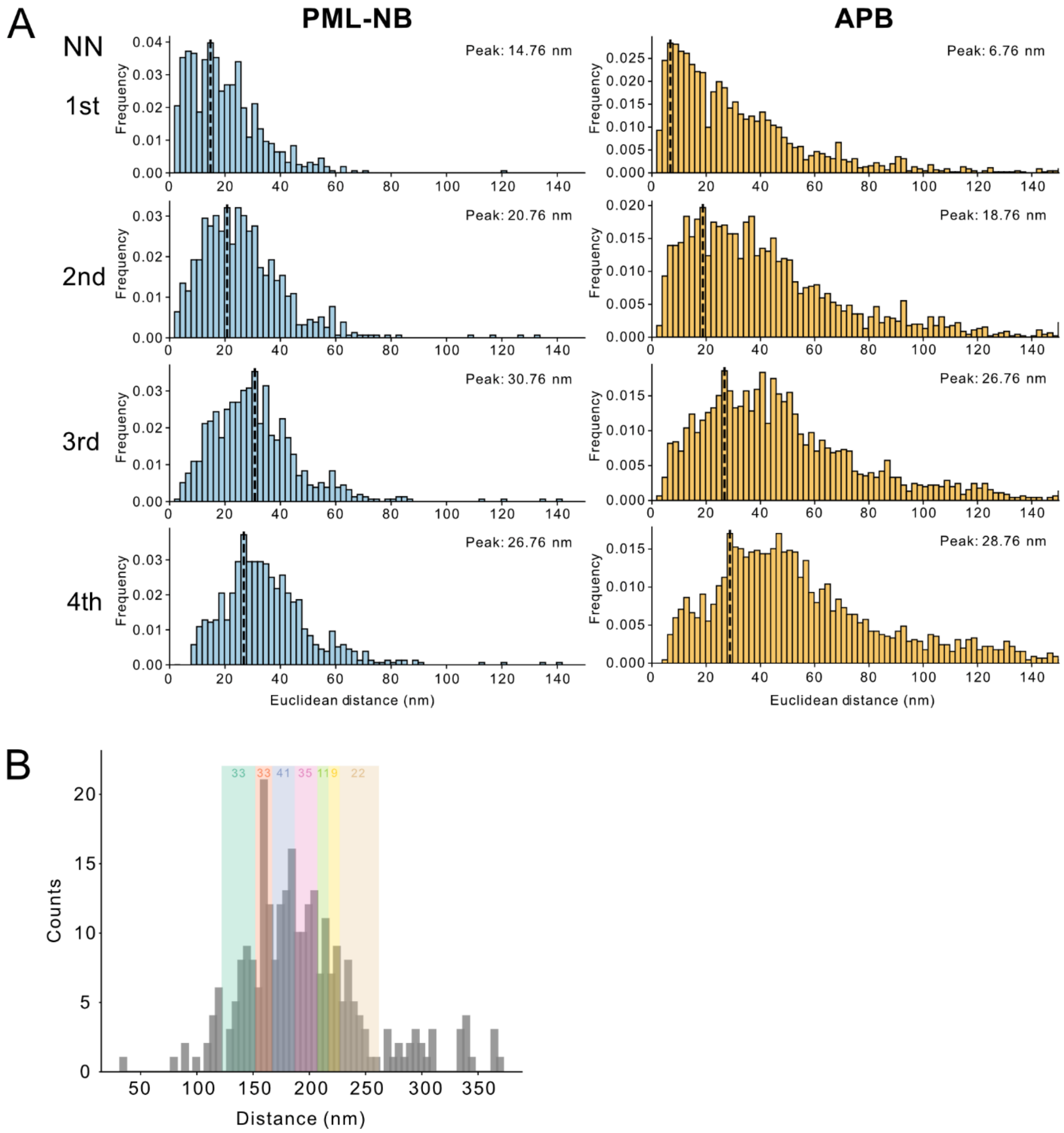


**Supplementary Figure 2. Nearest-neighbor distance analysis of PML molecules in APBs compared to canonical PML nuclear bodies (PML-NBs).** (**A**) Nearest-neighbor (NN) distance distributions for 3D Euclidean distances from each PML molecule to its 1st through 4th nearest neighboring molecules, shown separately for PML-NBs (blue) and APBs (orange). The highest peak distance for each NN order is indicated in nanometers. Histograms were computed using a 2 nm bin width. Data were collected across at least three independent replicates (n = 27 APB acquisitions, n = 10 PML-NB acquisitions). (**B**) Representative example of PML shell thickness calculation for a single APB. Local minima (valleys) in an inverted histogram of Euclidean distances from each PML molecule to the fitted sphere center (x-axis, in nm) were identified using SciPy's find_peaks function (colored blocks indicate identified valley positions). The first and last valley were assigned as the inner and outer radii of the PML shell, respectively, and shell thickness was calculated as the difference between these two values.

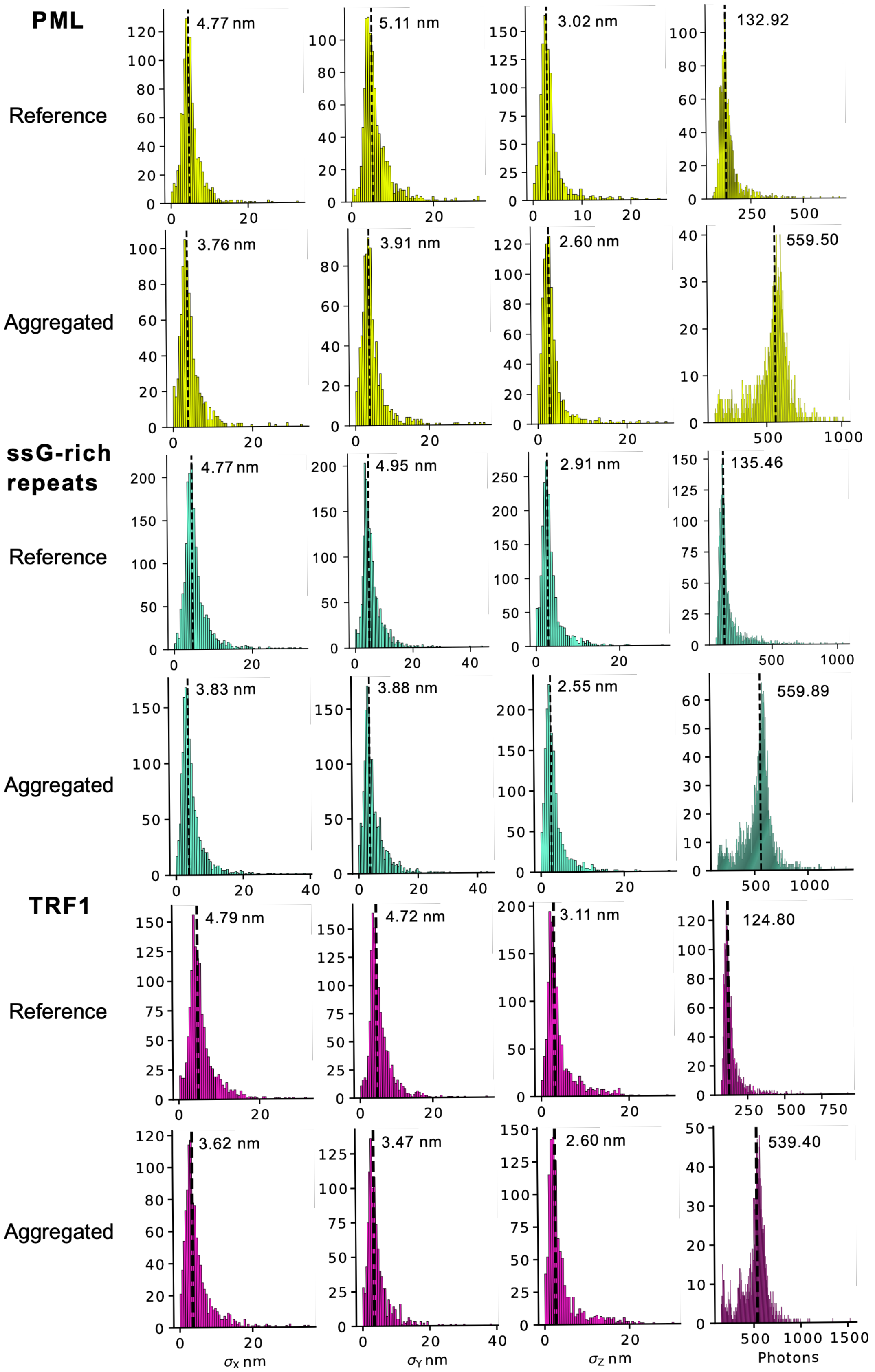


**Supplementary Figure 3. Photon count aggregation improves localization precision without affecting nearest-neighbor distances.** Comparison of per-trace localization precision distributions obtained with photon-aggregated data (500 photons per localization) and non-aggregated (raw) localizations for x, y and z dimensions. Vertical dashed lines indicate the median localization precision for each condition. Photon aggregation improved median localization precision by approximately 1 nm in x and y and 0.5 nm in z.

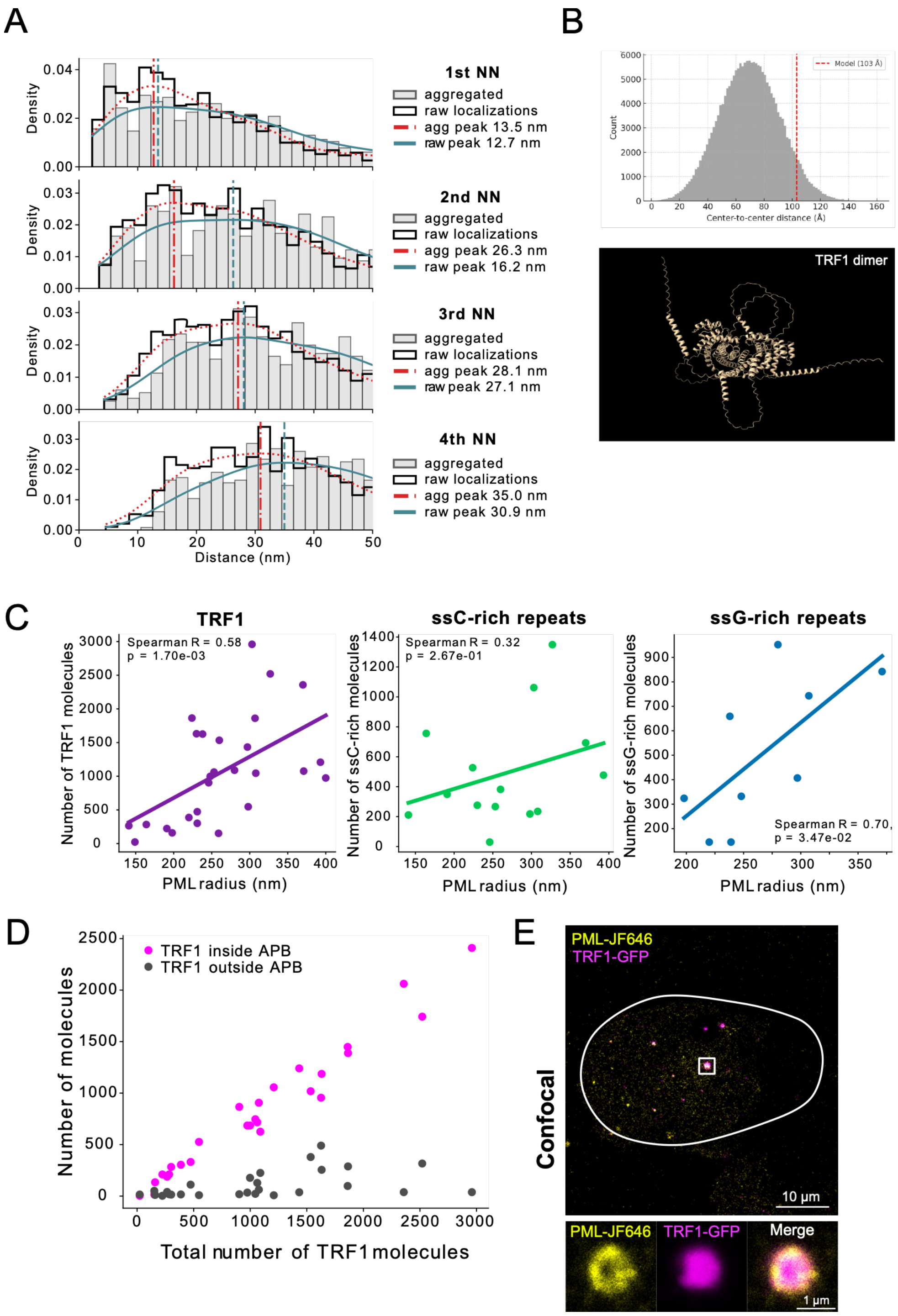


**Supplementary Figure 4. Quantification of APB component scaling and TRF1 organization.** (**A**) Kernel density estimation (KDE) of TRF1–TRF1 nearest-neighbor (NN) distances for aggregated and non-aggregated localizations. The first NN KDE peak position was not significantly altered by photon aggregation (12.7 nm versus 13.5 nm for non-aggregated and aggregated data, respectively), confirming that photon aggregation does not improve the measured molecular distances. (**B**) AlphaFold2 structural model of the TRF1 homodimer with flexible linkers and GFP fused at the C-terminus of each monomer, as used in the TRF1-GFP construct. A Monte Carlo simulation with two independent flexible linkers yielded an estimated center-to-center GFP distance of 7 ± 2 nm. The full-length TRF1-GFP dimer model is shown for reference. (**C**) Spearman correlation analysis of APB shell radius versus the number of detected molecules per APB for TRF1 (R = 0.58, magenta), ssG-rich repeats (R = 0.70, cyan) and ssC-rich repeats (R = 0.32, green), indicating that TRF1 and ssG-rich repeat abundance scale with APB size, whereas ssC-rich repeat abundance is less strongly associated with shell radius. (**D**) Quantification of TRF1 molecule counts for assemblies localized inside (pink) and outside (grey) the APB shell. The majority of TRF1 molecules localize inside the APB. TRF1 assembly sizes ranged from approximately 200 to 3000 molecules. (**E**) Representative confocal image confirming colocalization of HaloTag-PML detected with Janelia Fluor 646 ligand (yellow) and TRF1-GFP (magenta) in a U2OS ALT$^+$ cell, used to identify APB structures prior to MINFLUX imaging. These are the confocal images matching the MINFLUX structure in main figure 3C. Scale bar, 10 µm (overview) and 1 µm (inset).

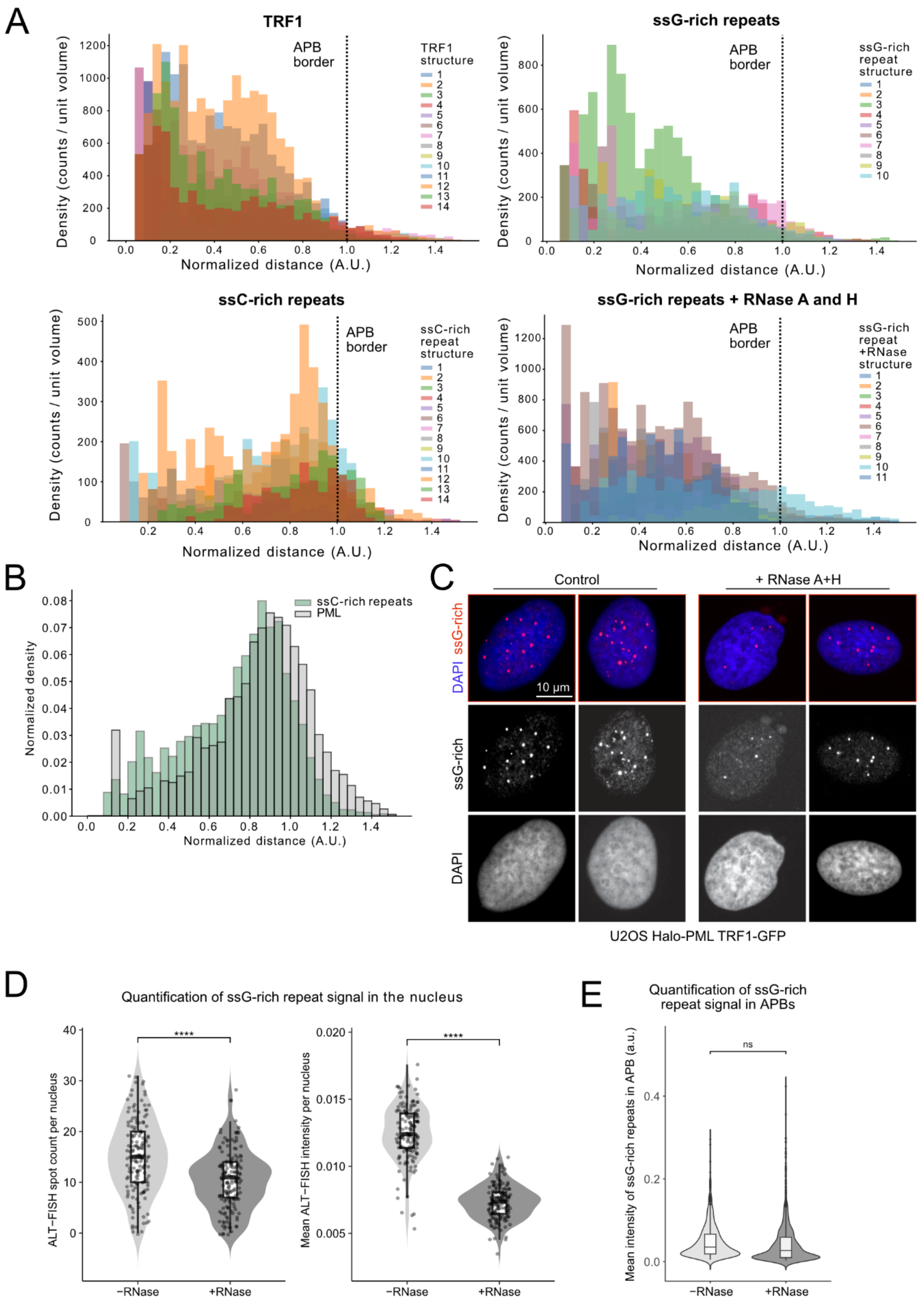


**Supplementary Figure 5. Radial distribution of APB components across individual acquisitions and validation of RNase treatment efficacy.** (**A**) Volume-normalized radial distance distributions for individual MINFLUX acquisitions of TRF1, ssG-rich repeats and ssC-rich repeats within APBs. Distances were calculated relative to the PML-defined APB center and normalized to the APB shell radius, such that the APB center corresponds to r = 0 and the shell boundary to r = 1. Individual histograms are shown to illustrate biological variability between acquisitions. Data were collected from at least three independent experiments for all conditions, except for ssG-rich repeats following RNase A and H treatment (two independent experiments). (**B**) Volume-normalized radial distance distributions of ssC-rich repeats (green) and PML molecules (grey) across all 3-target ssC-rich, PML and TRF1 acquisitions (n = 14). Distances are calculated relative to the PML-defined APB center and normalized to the APB shell radius, where r = 0 corresponds to the APB center and r = 1 to the shell boundary. The majority of ssC-rich repeat localizations are confined within the PML shell, with enrichment toward the inner shell surface. (**C**) Representative confocal fluorescence images of U2OS HaloTag-PML TRF1-GFP cells following RNase A and H treatment, showing the effect of RNA depletion on ssG-rich ALT-FISH signal. Scale bar, 10 µm. (**D**) Quantification of ssG-rich ALT-FISH signal in U2OS HaloTag-PML TRF1-GFP cells under control conditions and following RNase A and H treatment. Violin plots with overlaid boxplots and individual data points show ALT-FISH spot count (left) and mean ALT-FISH intensity per nucleus (right). ssG-rich ALT-FISH spot counts and signal intensity were significantly reduced upon RNase treatment (BH-adjusted Wilcoxon tests, p = 1.56e-7 and p = 4.14e-53, respectively), confirming successful RNA depletion. n = 177 nuclei for -RNase condition and 192 for +RNase condition. (**E**) Quantification of ssG-rich repeat signals inside APBs under control conditions and following RNase A and H treatment, assessed by confocal ALT-FISH imaging. RNase treatment resulted in a non-significant reduction of approximately 14% in ssG-rich signal intensity inside APBs (p = 0.058, n = 1024 APBs from 145 cells in control condition, 1576 APBs from 164 cells in RNase condition), indicating that the ssG-rich signal within APBs is predominantly single-stranded DNA.

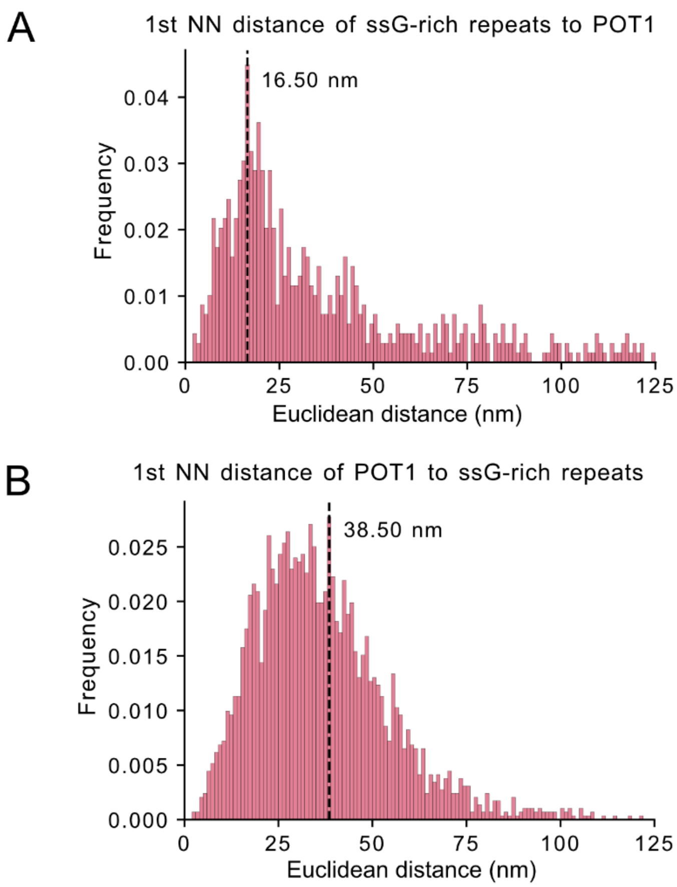


**Supplementary Figure 6. First nearest-neighbor (NN) distance analysis between POT1 and single-stranded (ss) G-rich repeats.** (**A**) Distribution of first NN distances from ssG-rich repeat localizations to the nearest POT1 localization. Distances were calculated from 3D MINFLUX data using Euclidean distance. The peak (mode) of the distribution is located at 16.5 nm. (**B**) Distribution of first nearest-neighbor distances from POT1 localizations to the nearest ssG-rich repeat localization. The peak (mode) of the distribution is located at 38.5 nm. Data were pooled from four independent acquisitions. The asymmetric NN distances reflect differences in spatial density and organization between POT1 and ssG-rich repeats.

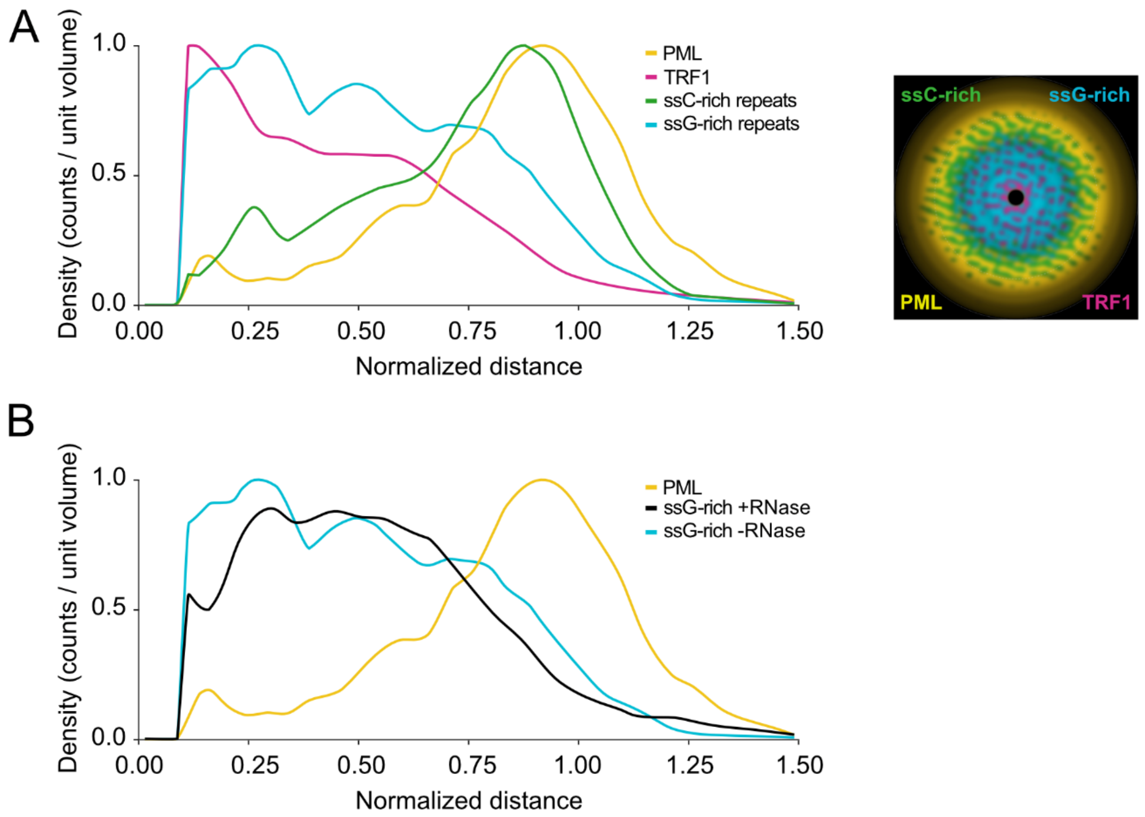


**Supplementary Figure 7. Radial abundance profiles reveal spatial segregation of components within APBs.** (**A**) Volume-normalized radial density profiles of PML (yellow), TRF1 (magenta), ssG-rich repeats (cyan) and ssC-rich repeats (green) across the APB, averaged across all acquisitions per condition. Each distribution is independently normalized to its maximum density to facilitate comparison of spatial enrichment patterns. TRF1 is enriched toward the APB center, ssG-rich repeats are broadly distributed across the interior, and ssC-rich repeats are enriched toward the inner PML shell surface. The right panel shows a schematic cross-section illustrating the distinct spatial compartmentalization of each component within the APB. (**B**) Comparison of volume-normalized radial density profiles of ssG-rich repeats under control conditions (cyan, n = 10) and following RNase A and H treatment (black, n = 11), shown together with the PML distribution (yellow) as a spatial reference. RNase treatment does not alter the radial distribution profile of ssG-rich repeats, consistent with the ssG-rich signal within APBs being predominantly DNA.

### Supplementary videos

**Supplementary Video 1. Three-dimensional MINFLUX rendering of a representative APB with three targets.** Individual localizations of PML (yellow), TRF1 (magenta) and ssG-rich telomeric repeats (cyan) are shown for the APB in Fig. 1E. The rendering performs one complete rotation about the vertical axis over 100 frames at 10 frames per second.

**Supplementary Video 2. PML channel of the APB shown in Supplementary Video 1.** Individual PML localizations (yellow) are shown with the same rotation and the same frames as the merged rendering.

**Supplementary Video 3. TRF1 channel of the APB shown in Supplementary Video 1.** Individual TRF1 localizations (magenta) are shown with the same rotation and the same frames as the merged rendering.

**Supplementary Video 4. ssG-rich telomeric repeat channel of the APB shown in Supplementary Video 1.** Individual ssG-rich repeat localizations (cyan) are shown with the same rotation and the same frames as the merged rendering.

**Supplementary Video 5. Three-dimensional MINFLUX rendering of a second representative APB with three targets.** Individual localizations of PML (yellow), TRF1 (magenta) and ssC-rich telomeric repeats (green) are shown for the APB in Fig. 4B. The rendering performs one complete rotation about the vertical axis over 100 frames at 10 frames per second.

**Supplementary Video 6. PML channel of the APB shown in Supplementary Video 5.** Individual PML localizations (yellow) are shown with the same rotation and the same frames as the merged rendering.

**Supplementary Video 7. TRF1 channel of the APB shown in Supplementary Video 5.** Individual TRF1 localizations (magenta) are shown with the same rotation and the same frames as the merged rendering.

**Supplementary Video 8. ssC-rich telomeric repeat channel of the APB shown in Supplementary Video 5.** Individual ssC-rich repeat localizations (green) are shown with the same rotation and the same frames as the merged rendering.
